# Modulation of ossification and inflammatory pathways during dexamethasone-induced *in vitro* osteogenesis

**DOI:** 10.64898/2026.02.12.705508

**Authors:** Antoine Buetti-Dinh, Claudia Siverino, Jorge Úbeda Garrido, Carmen Lanzillotti, Elisa Pianta, Gianvito Grasso, Sonja Häckel, Martin J. Stoddart, Elena Della Bella

## Abstract

**Background:** Dexamethasone (DEX) is used *in vitro* to promote osteogenic differentiation of human bone marrow mesenchymal stromal cells (hBMSCs). In clinical use, however, glucocorticoids induce osteoblast and osteocyte apoptosis while increasing osteoclast survival, leading overall to osteoporosis and high fracture risk. The overall impact of DEX on the differentiation of human progenitor cells remains contradictory and not fully understood, highlighting the need for further investigation using sequencing approaches as *in vitro* results will naturally influence further translational research.

**Methods:** hBMSCs were induced to osteogenic differentiation for 7 days using different concentrations of either DEX or the nonsteroidal glucocorticoid receptor agonist (+)-ZK216348. cDNA library preparation and RNA sequencing (RNAseq) were performed using Oxford Nanopore Technologies. Differentially expressed genes and pathways associated to the transactivation or transrepression activity of DEX were identified. Sequencing results were validated by qPCR, protein analysis, and with a functional assay on peripheral blood mononuclear cells to determine the overall effect of the BMSC supernatant.

**Results:** Hierarchical clustering of RNAseq data identified eight subclusters with shared regulatory patterns. Enrichment analysis revealed that both upregulated and downregulated genes are involved in ossification and extracellular matrix organization pathways. Several pro- and anti-inflammatory genes were differentially regulated. qPCR analysis validated the upregulation of *CXCL1*, *CXCL8*, *IL18*, and *COL8A1*, while *MMP1* and *CXCL12* expression decreased in response to DEX. Comparing DEX results with those obtained using (+)-ZK216348 helped distinguish the potential mechanisms regulating the expression of specific genes. Notably, *CXCL8* upregulation occurred through transactivation, whereas *COL8A1* upregulation is downstream of a transrepressed gene. Further *in vitro* experiments confirmed that DEX significantly increased *CXCL8* expression and IL-8 secretion. However, hPBMC responses indicated no significant pro- or anti-inflammatory effects from hBMSC conditioned medium.

**Conclusions:** In conclusion, the effects of DEX on the transcriptome of hBMSCs in a pro-osteogenic environment do not fully replicate the acquisition of an osteogenic phenotype. Several genes associated with ossification, extracellular matrix organization, and inflammation were dysregulated. The unique expression patterns of pro-inflammatory cytokines and collagen types warrant further investigation to elucidate their roles in osteogenic differentiation and bone homeostasis.

## 1. BACKGROUND

Understanding the mechanisms of bone formation is crucial for developing effective treatments for bone-related diseases. Researchers have used dexamethasone (DEX) as a tool to study osteogenesis in *in vitro* experimental models for more than thirty years, following early reports on its impact on the proliferation of bone cells in rodents and in osteogenesis of chick periosteum [1,2]. Since then, the use of DEX to induce osteogenic differentiation has become widespread and is now considered a standard protocol in the field [3–5]. DEX is used to promote mineralization *in vitro* and *in vivo*, and it is reported to function by increasing the expression and/or activity of the transcription factor RUNX2 [3,6–9]. However, using this drug to promote osteogenic differentiation contradicts its known clinical effects. Clinically, DEX promotes apoptosis in osteoblasts and osteocytes while increasing osteoclast survival, leading to osteoporosis, osteonecrosis, and a higher fracture risk [10].

Despite the clinical observations, DEX remains a standard tool in osteogenic differentiation protocols, with concentrations that are in the range of serum levels of patients treated with DEX [11]. It was previously demonstrated that DEX promotes osteogenic-like effects in human bone marrow mesenchymal stromal cells (hBMSCs) by downregulating *SOX9*, a key chondrogenic factor and a dominant negative of RUNX2 [12,13]. This downregulation should be sufficient to relieve inhibition on RUNX2, allowing differentiation to progress down the osteogenic lineage. However, the use of DEX simultaneously induces a strong upregulation in *PPARG* expression, leading to the parallel promotion of adipogenic differentiation in at least a subset of cells [12]. Overall, the effects of DEX on lineage commitment and differentiation of human progenitor cells remain contradictory and not completely understood.

DEX is a synthetic fluorinated glucocorticoid derived from cortisol, with a longer half-life and higher affinity to the glucocorticoid receptor (GR) compared to cortisol [14]. Upon binding to GR, the complex translocates to the nucleus where it controls the expression of target genes by either transactivation or transrepression [15]. Transrepression is mostly associated with the anti-inflammatory effects of DEX, as it blocks the expression of NF-kB target genes, while transactivation is usually associated with side effects [16]. In an attempt to dissociate these effects and improve the therapeutic profile of glucocorticoids, molecules inducing only transrepression of target genes were designed, such as the nonsteroidal GR agonists (+)-ZK216348 and AZD9567 [16,17]. In addition to their clinical potential, these molecules serve as invaluable tools *in vitro* to determine whether the effects of glucocorticoids are mediated through transactivation or transrepression. This approach helps clarify the molecular cascade following glucocorticoid stimulation.

e aim of this study was to investigate the broader effects of DEX on the differentiation of human BMSCs under pro-osteogenic conditions. To this end, bulk RNA sequencing (RNA-seq) was employed to profile the transcriptomic response of hBMSCs to varying concentrations of DEX or the selective glucocorticoid receptor modulator (+)-ZK216348. Differentially expressed genes (DEGs) identified by RNA-seq were subsequently validated in independent experiments through gene and protein expression analyses. Additionally, a functional assay using peripheral blood mononuclear cells (PBMCs) was conducted to assess the immunomodulatory effects of BMSC-conditioned media.

## 2. MATERIALS AND METHODS

### 2.1 Overall experimental design

The study was conducted in three main phases.

- Transcriptomic profiling and initial validation. Previously collected osteogenic samples treated with different concentrations of DEX or (+)-ZK216348 were used for RNA-seq analysis. Selected DEGs were validated at the gene expression level.

- Independent validation experiment. A new osteogenic differentiation experiment was performed to further validate key findings at both the transcript and protein levels.

- Functional assay on PBMCs. Conditioned media from treated BMSCs were applied to PBMC cultures to evaluate the net effect on inflammatory responses.

Figure 1 provides a graphical summary of the experimental workflow. Full details are provided in the following paragraphs.

**Fig. 1.**
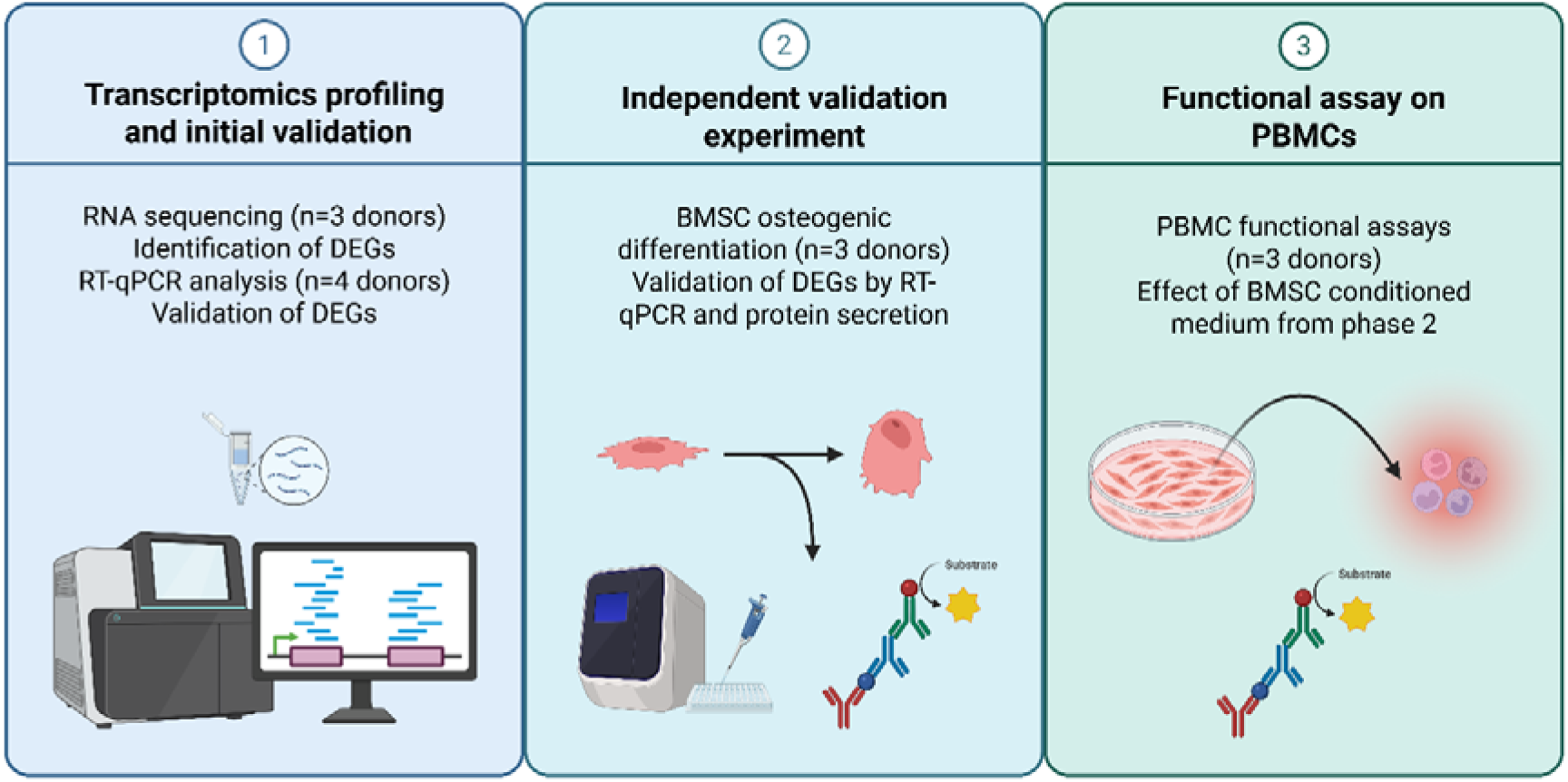
Graphical summary of the experimental workflow. The study was conducted in three sequential phases. 1) Transcriptomic profiling and initial validation, that allowed the identification of DEGs following DEX or (+)-ZK216348 in pro-osteogenic conditions. 2) Further validation of results in an independent experiment. 3) Functional assay on PBMCs to evaluate the immunomodulatory effects of BMSC-conditioned media. RT-qPCR: Reverse Transcription quantitative Polymerase Chain Reaction. DEG: Differentially Expressed Gene. BMSC: Bone Marrow-derived Mesenchymal Stromal Cells. PBMC: Peripheral Blood Mononuclear Cells. Created with BioRender.com.

### 2.2 Biological materials used for RNAseq

In the first phase of the study, the RNA and cDNA samples generated for a previous study [12] were used for initial RNAseq and qPCR validation of sequencing results. In brief, hBMSCs were induced to osteogenic differentiation using either 0, 10, or 100 nM DEX-cyclodextrin complex (water-soluble formulation), or using the same concentrations of (+)-ZK216348 as a comparison to evaluate whether DEX effects are induced via a transactivation or a transrepression mechanism. Samples were collected for total RNA isolation and cDNA synthesis. Samples remaining from this study were stored at −80°C (RNA) or −20°C (cDNA) until further analysis. For full experimental details refer to Della Bella *et al.* [12].

For the purposes of the present study, the samples from *n*=3 donors (M, 24y; M, 48y; F, 45y) were selected for RNAseq. In summary, the samples from each donor included the following groups:

- **CRL**: cells in growth medium (Dulbecco’s Modified Eagle Medium–DMEM 1 g/l glucose, 10% Fetal Bovine Serum–FBS, 100 U/mL penicillin, 100 μg/mL streptomycin).
- **AA-GP**: cells in osteogenic basal medium (growth medium + 50 μg/ml L-Ascorbic Acid 2-phosphate sesquimagnesium salt hydrate + 5 mM β-Glycerol Phosphate).
- **DEX10**: cells in osteogenic basal medium + 10 nM DEX.
- **DEX100**: cells in osteogenic basal medium + 100 nM DEX.
- **ZK10**: cells in osteogenic basal medium + 10 nM (+)-ZK216348.
- **ZK100**: cells in osteogenic basal medium + 100 nM (+)-ZK216348.
- **DMSO10** (vehicle control): cells in osteogenic basal medium + DMSO (same dilution as that used for 10 nM (+)-ZK216348).
- **DMSO100** (vehicle control), cells in osteogenic basal medium + DMSO (same dilution as that used for 100 nM (+)-ZK216348).

The selected RNA samples were subjected to a DNase treatment to remove contaminant genomic DNA. For validation of RNAseq results, the cDNA previously synthesized [12] was used for qPCR analysis of the target genes, using the same donors used for sequencing and an additional one from the previous study (F, 37y). To keep costs manageable, only one technical replicate per group and donor was used for sequencing. qPCR validation was performed in technical duplicates for each group and donor.

### 2.3 RNA sequencing and bioinformatics

DNase-treated total RNA (50 ng in a 5 μl volume) was used for Oxford Nanopore Technologies (ONT) library preparation according to manufacturer instructions (Kit SQK-PCB109) and sequencing was performed with an ONT R9.4 flow cell. Accurate basecalling was performed with ONT Guppy basecalling software version 5.0.7+2332e8d65 (configuration file "\dna r9.4.1 450bps hac.cfg") followed by barcoding with ONT Guppy barcoding software version 5.0.11+2b6db_a5. ONT Pychopper (v2) was then used to assess RNA quality and process sequencing reads for further analysis. Finally, the ONT long-reads pipeline for differential gene expression (DGE) and differential transcript usage (DTU) analysis [18] was used for transcripts quantification. The snakemake computational pipeline included minimap2, salmon, edgeR, DEXSeq and stageR. The transcriptome ("\Homo sapiens.GRCh38.cdna.all.fa.gz") and annotation ("\Homo sapiens.GRCh38.95.gtf.gz") files were downloaded from the ftp.ensembl.org repository. Default options for the pipeline were used: minimap index opts: ""; minimap2 opts: ""; maximum secondary: 100; secondary score ratio: 1.0; salmon libtype: "A"; min samps gene expr: 3; min samps feature expr: 1; min gene expr: 10; min feature expr: 3.

RNAseq results were filtered by false discovery rate (FDR)<0.05 and Log2FC>|1|. Principal component analysis (PCA) and heatmaps were applied to the scaled date using the "prcomp" and "heatmap.2" functions of the R language. Clustering in heatmaps used average linkage and Pearson distance. Functional categories were assigned to the gene clusters by performing a Gene Ontology (GO) Enrichment Analysis using the clusterProfiler package in R [19–21].

### 2.4 Osteogenic differentiation of hBMSCs

To validate sequencing data in an independent set of experiments, hBMSCs (*n*=3; M, 18y; M, 18y; M, 31y) were isolated and expanded as previously described [12]. Bone marrow was collected from patients undergoing spine surgery at the Inselspital Bern with signed informed patient consent. The Swiss Human Research Act does not apply to research that utilizes anonymized biological material and/or anonymously collected or anonymized health-related data. Therefore, this project did not need to be approved by an ethics committee. Patients’ general consent was obtained, which also covers the anonymization of health-related data and biological material. The study was conducted in accordance with the Declaration of Helsinki.

hBMSCs at passage 3 were seeded in triplicate into 12-well plates at a density of 1.5 × 10_ cells/cm² and incubated overnight in expansion medium (α-MEM supplemented with 10% FBS, 100 U/mL penicillin, and 100 μg/mL streptomycin). After cell adhesion, osteogenic differentiation was induced using medium containing DMEM (1 g/L glucose), 10% FBS, 1% Pen/Strep, 50 μg/mL L-ascorbic acid 2-phosphate sesquimagnesium salt hydrate, and 5 mM β-glycerophosphate, supplemented with 0, 10, or 100 nM DEX-cyclodextrin complex. Undifferentiated controls were maintained in growth medium (DMEM 1 g/L glucose, 10% FBS, 1% Pen/Strep). Medium was refreshed three times per week, and cells were cultured for up to 21 days.

Conditioned medium and RNA samples were collected on day 7, while mineral deposition was assessed by Alizarin Red staining on day 21. For each donor, conditioned medium was pooled from multiple wells, yielding 6 mL per condition, collected under sterile conditions and stored at −20°C until analysis.

### 2.5 Alizarin Red staining and quantification

Alizarin Red staining was used to assess mineral deposition as previously described [12]. After fixation with 4% neutral buffered formalin, monolayers were stained with 40 mM Alizarin Red S solution, pH 4.2 (Sigma-Aldrich). Images were captured after extensive washing using an inverted microscope (Axio Vert.A1, Carl Zeiss AG, Oberkochen, Germany). Semi-quantitative analysis was performed after elution of bound Alizarin Red using the cetylpyridinium chloride method [12]. Absorbance was measured at 540 nm using an Infinite® 200 PRO microplate reader (Tecan, Männedorf, Switzerland).

### 2.6 RNA isolation and reverse transcription

RNA was isolated from cells at day 7 using a standard phenol-chloroform extraction as previously described [12]. Quantity and purity of RNA were evaluated using a Nanodrop One UV-Vis spectrophotometer (Thermo Fisher). One microgram of total RNA was reverse transcribed using the TaqMan Reverse Transcription reagents (Applied Biosystems, Foster City, CA, USA), following manufacturer’s instruction.

### 2.7 Gene expression analysis

For qPCR validation of RNAseq targets, cDNA was obtained from both the original sample set used in in the first phase of the study [12] and the additional cohort of donors from the second phase. 10 ng of cDNA were used for qPCR using the TaqMan Gene Expression Master Mix (Applied Biosystems) in a QuantStudio 7 Flex Real-Time PCR system (Applied Biosystems). Assay details are reported in **Table 1**.

**Table 1.**
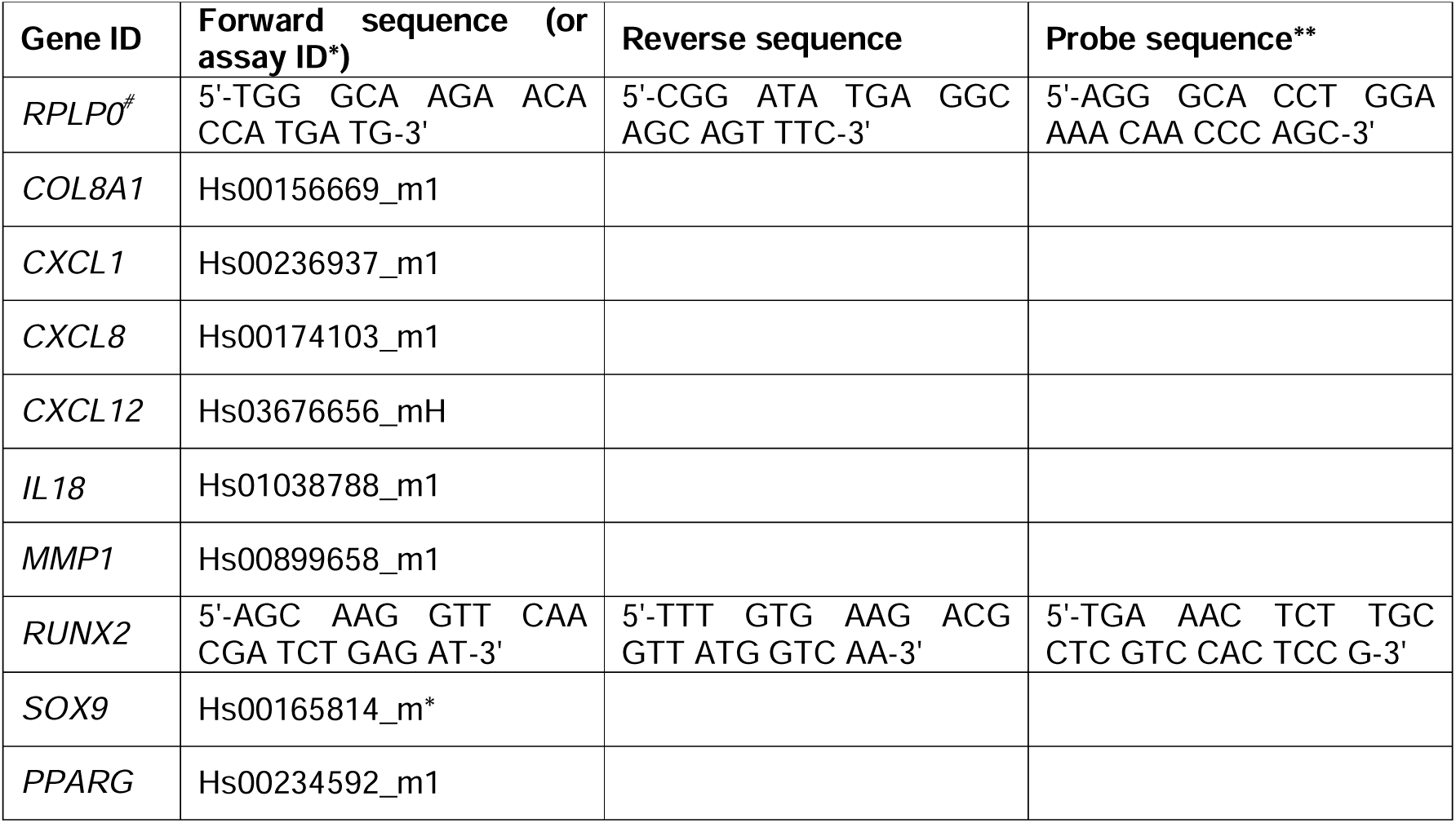
qPCR primers details. * TaqMan Gene expression assay (Thermo Fisher). 5’ modification: FAM. 3’ modification: NFQ-MGB. ** 5’ modification: FAM. 3’ modification: TAMRA. # Reference gene.

### 2.8 Validation of IL-8 and IL-18 expression by ELISA

The concentration of secreted IL-8 and IL-18 in hBMSC conditioned medium were measured using enzyme-linked immunosorbent assay (ELISA) kits (DuoSet, R&D Systems, Minneapolis, MN, USA). The ELISA was performed according to the manufacturer’s instructions, and absorbance was measured at 450 nm. Concentrations were determined by comparing absorbance values to a standard curve generated with known concentrations of recombinant IL-8 and IL-18. Unconditioned medium (*i.e.*, medium that was not in contact with cells) was tested by ELISA to exclude background signal from serum components.

### 2.9 Effect of cytokine release on human peripheral blood mononuclear cells (hPBMCs)

hPBMCs were isolated from healthy human donors (*n*=3). The buffy coat was obtained from the Regionaler Blutspendedienst SRK Graubünden, and PBMCs were isolated via density gradient centrifugation using Histopaque®-1077 (Sigma-Aldrich, Merck KGaA, Darmstadt, Germany) at 800 g for 20_min at room temperature. PBMCs were treated for 24 h with the conditioned medium of BMSCs induced to osteogenic differentiation. Briefly, PBMCs were seeded at a density of 2×10^5^ cells/well in a 96 well plate in RPMI 1640 (containing L-Glutamine and HEPES) supplemented with 10% FBS and 1% Pen/Strep. 24 h after seeding, the plates were centrifuged to allow cell deposition, and medium was replaced with 100 μl fresh growth medium + 100 μl BMSC conditioned medium. Freshly prepared osteogenic medium containing the different concentrations of DEX were also used on PBMCs as a control. After an additional 24 h, samples were collected, cells were removed by centrifugation, and conditioned medium (both from BMSCs and from PBMCs) was assessed for IL-6, TNFα, IL-1β, IL-12p70, and IL-10 production using a V-plex multiplex assay (MSD, Rockville, MD, USA).

PBMCs from the same donors were stained with 1.5 μM PKH-26 (Sigma-Aldrich) and seeded in 12-well plates at a density of 0.5×10^6^ cells/well, in 600 μl RPMI 1640, 10% FBS, 1% Pen/Strep. Twenty-four hours after seeding, 600 μl of BMSC conditioned media were added. PBMCs were incubated for 72 h at 37°C, 5% CO_2_, 90% rH, then analysed by flow cytometry (FACSAria III, Becton Dickinson, Franklin Lakes, NJ, USA). Proliferation analysis was performed using the Proliferation Tool on FlowJo v.10.10.0 (Becton Dickinson) and the percentage of divided cells was reported.

### 2.10 Statistical analysis and data visualization

GraphPad Prism 10 for Windows (GraphPad Software, San Diego, CA, USA) was used for data visualization and statistical analysis. For both qPCR and cytokine measurements, a two-way ANOVA was performed with treatment and donor as independent variables to assess their effects on gene expression and cytokine levels. Dunnett’s multiple comparisons test was applied to evaluate differences between treatment groups relative to the control, focusing on the main effect of treatment. Sidak Differences were considered statistically significant when *p* < 0.05.

## 3. RESULTS

### 3.1 Transcriptomic profiling and initial validation reveals modulation of ossification, extracellular matrix, and inflammatory mediators with mixed effects

Principal component analysis (PCA) revealed that hBMSCs treated with increasing concentrations of DEX exhibited progressive transcriptomic changes. At 10 nM, DEX-treated samples clustered closely with untreated controls (AA-GP and DMSO), while 100 nM DEX caused a distinct shift along PC1, indicating stronger transcriptional modulation. The selective GR agonist (+)-ZK216348 also influenced gene expression, but to a lesser extent than DEX, particularly at lower concentrations (Fig. 2A). Hierarchical clustering of RNAseq data identified eight subclusters of genes with shared regulatory patterns across treatment conditions. Subcluster 3 included genes downregulated by DEX, while subclusters 4 and 5 comprised genes upregulated by DEX. These clusters reflect the dual regulatory role of DEX in modulating gene expression under osteogenic conditions (Fig. 2B). A more detailed version of the heatmap in Fig. 2B, with additional details about the subclusters, can be found in the **Supplementary File 1**.

**Fig. 2.**
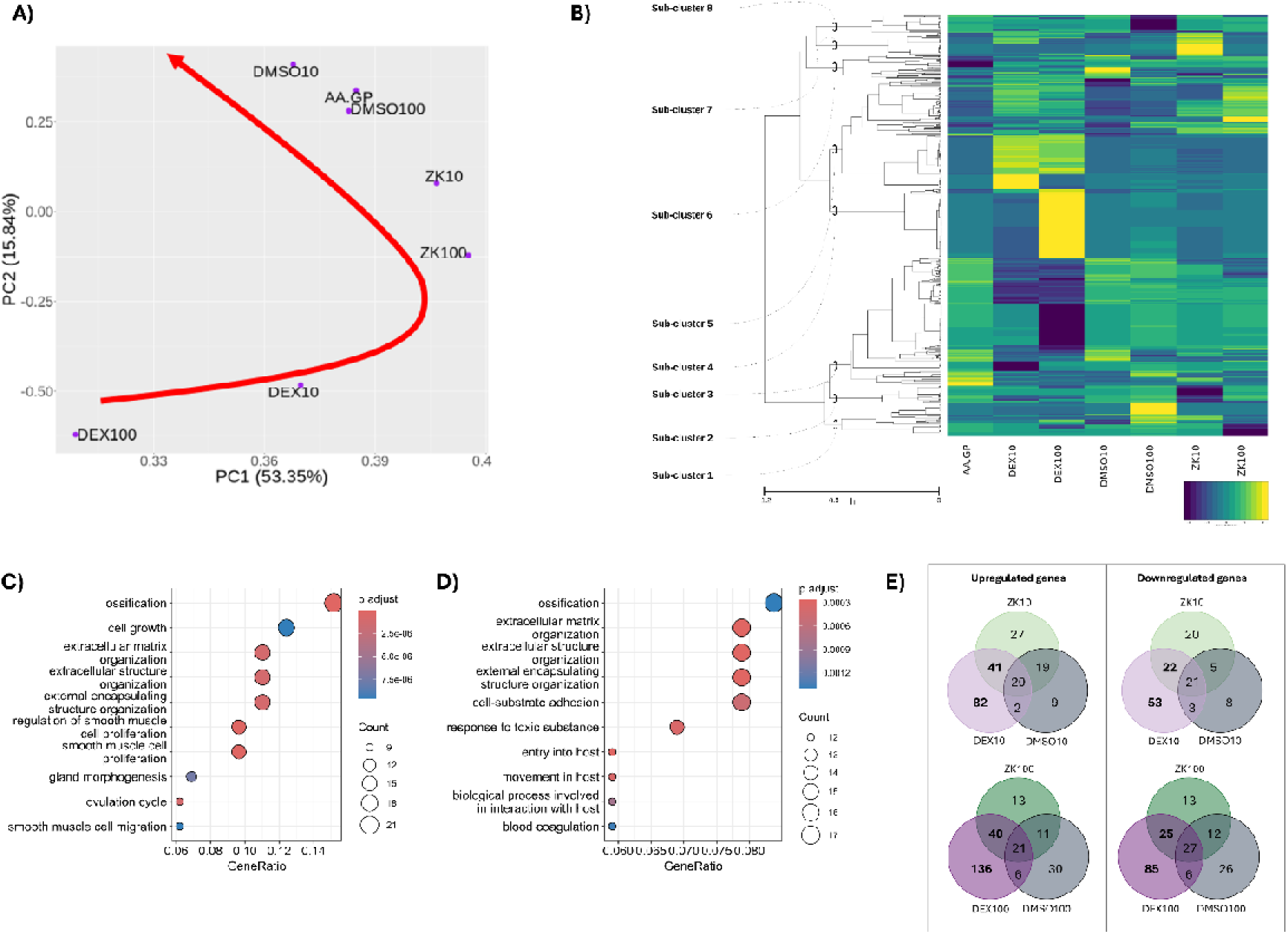
Transcriptomic profile of hBMSCs identified by RNA sequencing (n=3 biological replicates). **A)** Principal Component Analysis. Loadings are the chemical inducers at different concentrations. The red arrow indicates increasing differences in the number of differentially regulated genes. **B)** Heatmap and hierarchical clustering of genes identified by RNAseq based on their expression profile over the different experimental conditions (represented on the x-axis, in which DEX: dexamethasone; ZK: (+)-ZK216348, DMSO: dimethyl sulfoxide, AA-GP: Osteogenic basal control; the number indicates concentrations in nM). Heatmap colour ranges from dark blue (less expressed) to yellow (more expressed). **C)** Gene Ontology (GO) Enrichment analysis for genes downregulated by DEX (subcluster 3). **D)** GO Enrichment analysis for genes upregulated by DEX (subclusters 4 and 5). **E)** Venn diagrams to identify genes that are regulated by dexamethasone only (indicating transactivation) or by both DEX and (+)-ZK216348 (indicating involvement of transrepression).

Gene Ontology (GO) enrichment analysis showed that both upregulated and downregulated genes were significantly associated with ossification and extracellular matrix organization (Fig. 2C**-D**). DEX treatment resulted in the upregulation of several pro-inflammatory cytokines, including *CXCL1*, *CXCL8*, and *IL18*, while simultaneously downregulating others such as *IL6* and *CXCL12*. This mixed regulatory pattern highlights the complex role of DEX in modulating inflammation during osteogenic differentiation. **Table 2** lists the genes identified by GO analysis in these pathways. In Fig. 2E, Venn diagrams of upregulated and downregulated genes allowed the identification of genes which are uniquely regulated by DEX, (+)-ZK216348, or DMSO, or commonly regulated.

**Table 2:** List of genes identified by Gene Ontology enrichment analysis related to GO:0001503 (Ossification) and GO:0030198 (Extracellular matrix organization) in subclusters 3 and 4+5.

| Subcluster 3<br>Genes downregulated by DEX |  | Subcluster 4+5<br>Genes upregulated by DEX |  |
| --- | --- | --- | --- |
| Ossification<br>(GO:0001503) | Extracellular matrix<br>organization<br>(GO:0030198) | Ossification<br>(GO:0001503) | Extracellular matrix<br>organization<br>(GO:0030198) |
| ADAMTS12 | ADAMTS1 | ADAMTS7 | ADAMTS7 |
| ANKH | ADAMTS12 | ATP2B1 | ANXA2 |
| CDH11 | COL10A1 | CLEC3B | CAV2 |
| CTSK | COL12A1 | COL11A1 | COL11A1 |
| DDR2 | COL14A1 | CTNNB1 | COL8A1 |
| DHRS3 | CTSK | GPM6B | CST3 |
| IGFBP3 | CYP1B1 | HDAC7 | FBLN1 |
| IGFBP5 | DDR2 | IGF2 | FBLN5 |
| IL6 | IL6 | LEP | GPM6B |
| ISG15 | LUM | MMP13 | LAMA2 |
| ITGA11 | MMP1 | PRKACA | LOXL3 |
| MDK | MMP2 | RANBP3L | MMP13 |
| MMP2 | POSTN | SCUBE3 | MMP7 |
| PENK | SULF1 | SORT1 | OLFML2A |
| PTN |  | SRGN | PTX3 |
| TNC |  | ZBTB16 | VIT |
| TNFAIP6 |  |  |  |
| VCAN |  |  |  |

The heatmap in Fig. 2B and has been re-ordered based on the levels of DEX response (see **Fig. S2** and associated data in **Supplementary file 1**). On top of DEX-sensitive, positively stimulated genes, the acute phase *SAA1* gene (Serum Amyloid A1) can be found, together with tissue remodelling genes including *CADM3* (Cell Adhesion Molecule 3), *TIMP4* (Metalloproteinase inhibitor 4), *MMP7* (Matrix Metallopeptidase 7), *MMP13* (Matrix Metallopeptidase 13), and *OMD* (Osteomodulin). *CXCL1* (C-X-C Motif Chemokine Ligand 1), *CXCL8* (C-X-C Motif Chemokine Ligand 8), and *IL18* (Interleukin 18) were also found close to the top of the highly positively DEX-regulated genes, despite their proinflammatory role. Conversely, DEX-sensitive, negatively regulated genes comprised *SFRP4* (Secreted Frizzled Related Protein 4), as well as other genes involved in tissue remodelling, such as *MMP1* (Matrix Metallopeptidase 1), *COL10A1* and *COL14A1* (Collagen Type X Alpha 1 Chain and Collagen Type XIV Alpha 1 Chain, respectively), *KRT16* (Keratin 16), *HAPLN1*/3 (Hyaluronan And Proteoglycan Link Protein 1 and 3), *FNDC1* (Fibronectin Type III Domain Containing 1), and *JUP* (Junction Plakoglobin). Genes involved in immune system regulation, such as *IL6*, were also found ranked near the top of strongly negatively DEX-regulated genes.

Comparison of DEX results with the selective GR agonist (+)-ZK216348 enabled the differentiation between transactivation and transrepression mechanisms. Genes uniquely regulated by DEX were attributed to transactivation, while those shared between DEX and ZK were considered transrepression targets. Venn diagram representations in Fig. 2E were used to identify genes regulated by transactivation (DEX unique) *vs.* transrepression (DEX/ZK shared). **Supplementary file 2** contains information about all the genes which are comprised within the DEX unique and the DEX/ZK shared groups. Considering the different IC_50_ of DEX and (+)-ZK216348 on the glucocorticoid receptors (respectively ∼5 and ∼20 nM), the genes which are uniquely regulated by DEX at the 10 nM dose but become DEX/ZK shared at 100 nM are here considered to be downstream targets of GR transrepression (see “IC_50_ effect” tab in **Supplementary File 2**).

For the genes whose expression is controlled by transactivation, 46 upregulated genes overlap between the DEX10 and DEX100 unique genes, and 21 are downregulated both by DEX10 and DEX100 only. For most of those genes, there is a dose-dependent trend on the log2FC (see “DEX only Dose-dependency” tab in **Supplementary File 2**).

Of note, *CXCL1*, *CXCL8*, and *IL18* were upregulated via transactivation, whereas *MMP1*, *CXCL12*, and COL8A1 appeared to be regulated by transrepression. Dose-dependent trends further supported these regulatory distinctions. These genes were selected for further validation based on both biological relevance and statistical significance (**Table 3**).

**Table 3:**
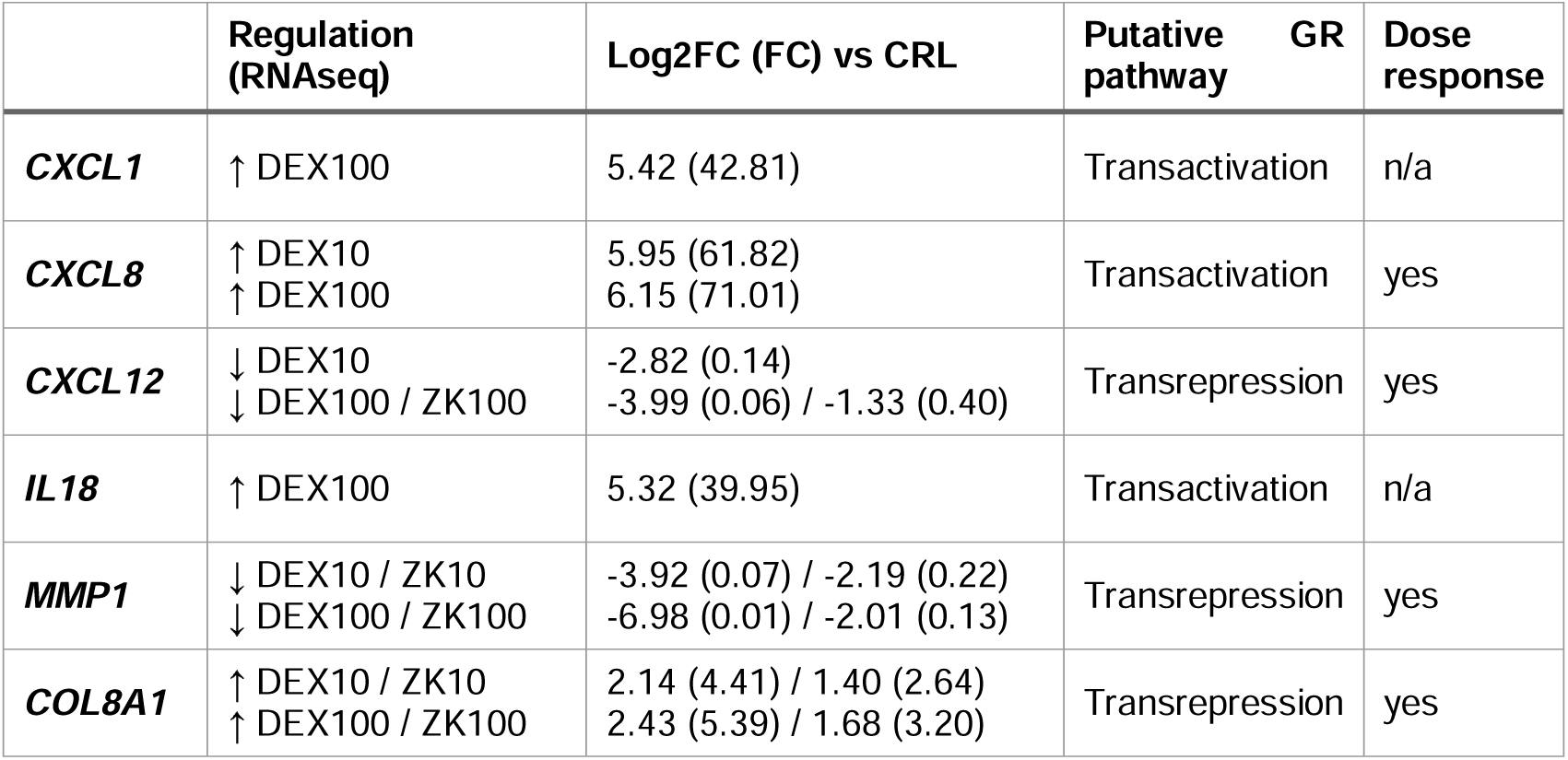
Summary of targets selected for qPCR validation. FC: Fold Change. GR: Glucocorticoid Receptor. DEX: Dexamethasone.

qPCR analysis confirmed the RNAseq results, showing significant upregulation of *CXCL1*, *CXCL8*, *IL18*, and *COL8A1* in response to DEX treatment compared to osteogenic basal medium (Fig. 3). Although (+)-ZK216348 showed a similar trend for these genes, the changes were not statistically significant. Conversely, *MMP1* and *CXCL12* were significantly downregulated by both DEX and ZK (Fig. 3), supporting their regulation via transrepression. These findings reinforce the differential regulatory mechanisms of DEX and validate the transcriptomic data.

**Fig. 3.**
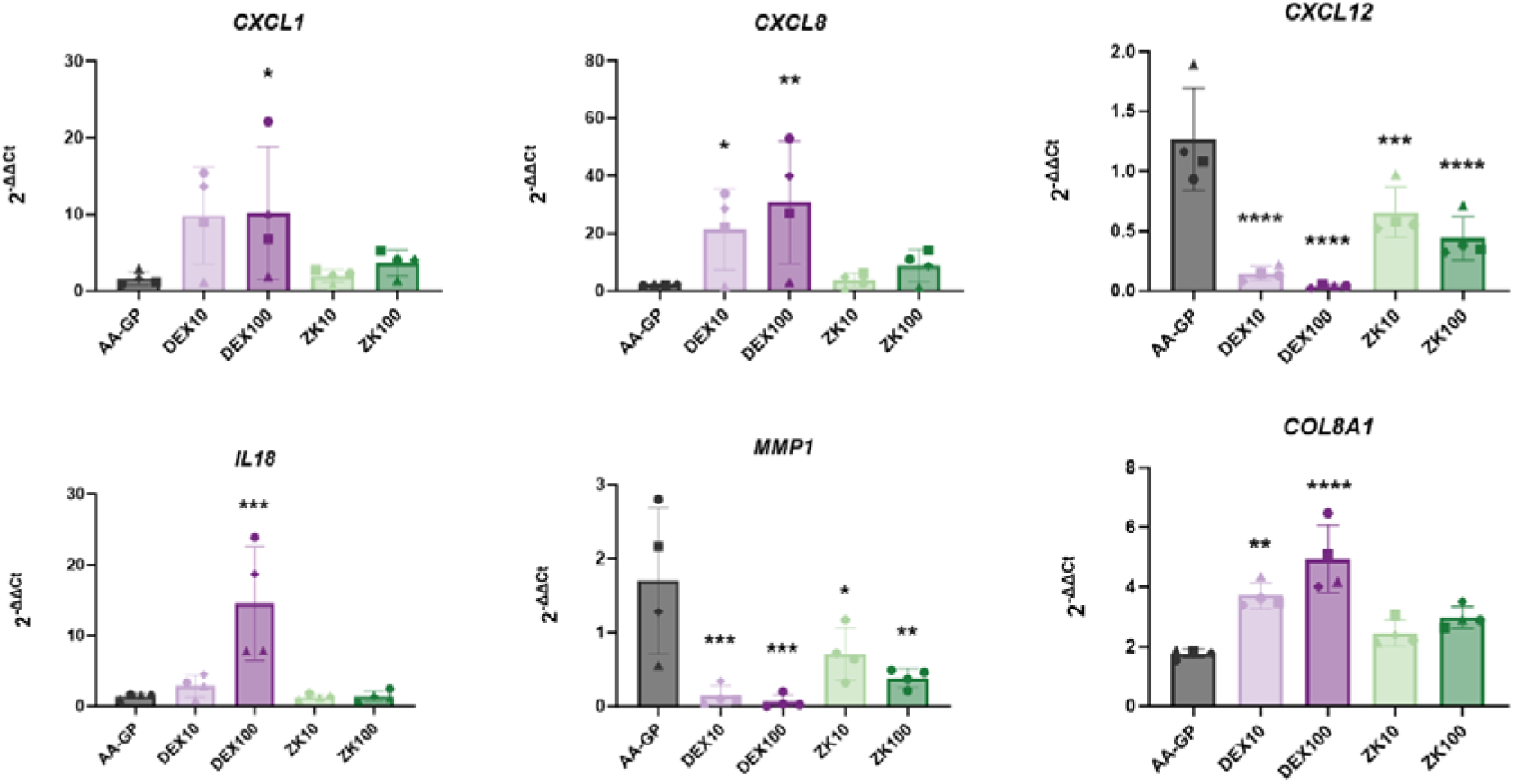
Validation of differentially expressed gene expression. CXCL1, CXCL8, CXCL12, IL18, MMP1, and COL8A1 expression was validated by RT-qPCR. Relative gene expression was calculated as 2^-ΔΔCt^, with RPLP0 as endogenous control and the undifferentiated control as calibrator. * p<0.05; ** p<0.01; *** p<0.001; **** p<0.0001. A two-way ANOVA was used followed by a Dunnett’s test to compare the mean of each group to the AA-GP control. Each data point represents an individual donor (n = 4 biological replicates), with values shown as the average of two technical replicates (i.e., two separate wells per donor). The bars indicate the mean value across all donors, and the error bars show the standard deviation.

Overall, these transcriptomic insights prompted further validation in independent donor cohorts, as described below.

### 3.2 RNAseq data are validated in independent *in vitro* experiments

RNAseq data were further validated in independent *in vitro* experiments. hBMSCs from 3 donors underwent osteogenic differentiation as described above for groups CRL, AA-GP, DEX10, and DEX100. Gene expression analysis at day 7 and ARS at day 21 of BMSCs was performed to further validate previously observed trends.

As expected, ARS staining increased with increasing DEX concentrations (Fig. 4A), together with *SOX9* downregulation and *PPARG* upregulation (Fig. 4B). In this set of donors, *RUNX2* levels were reduced in the presence of 10 nM DEX (Fig. 4B).

**Fig. 4.**
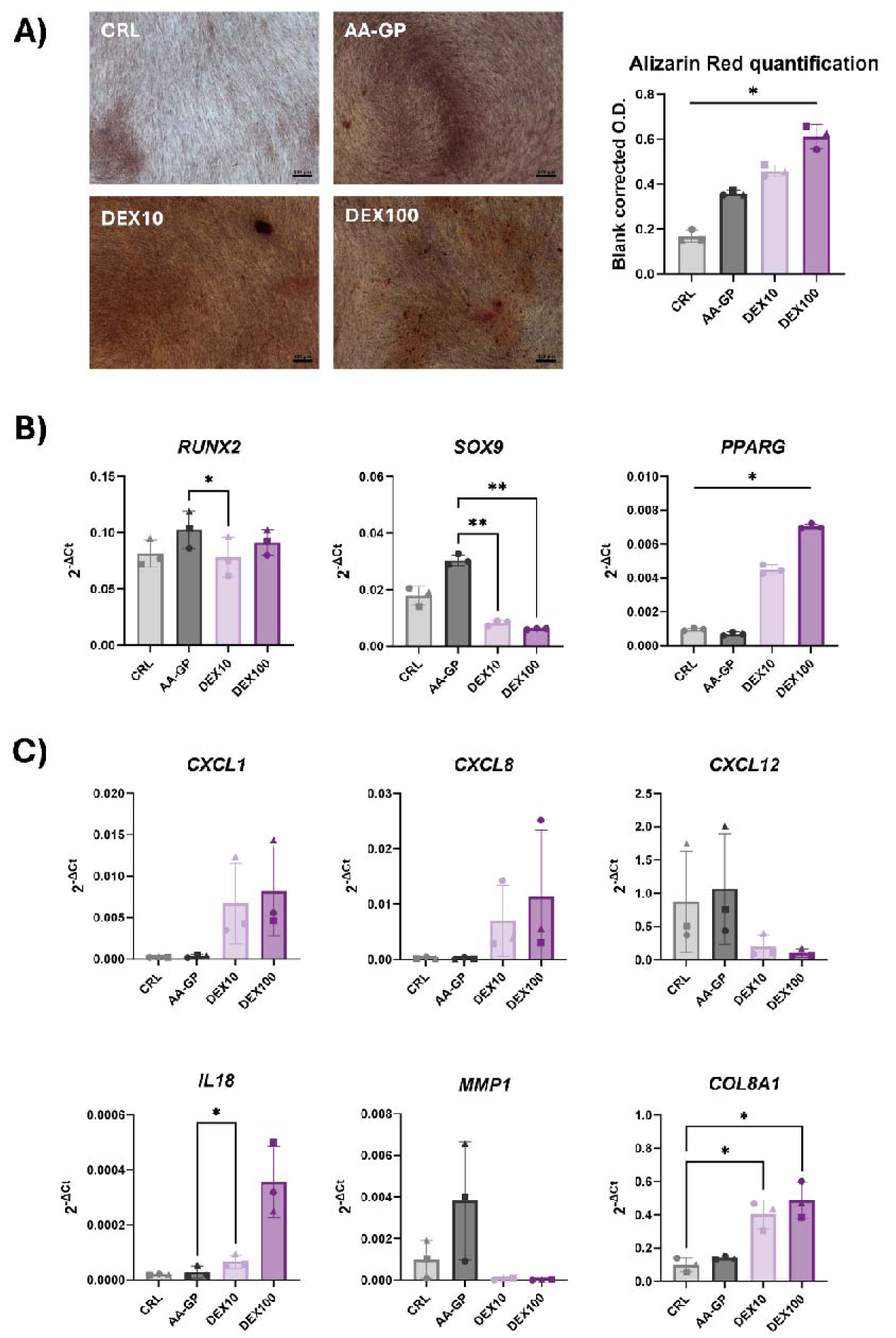
Validation of RNA sequencing results in a new cohort of donors. **A**) Alizarin Red representative pictures and quantification. **B**) Expression of trilineage differentiation markers RUNX2, SOX9, and PPARG. **C**) Expression of genes identified by RNAseq as differentially expressed. * p<0.05; ** p<0.01. A two-way ANOVA was used followed by a Dunnett’s test to compare the mean of each group to the CRL group. Each data point represents an individual donor (n = 3 biological replicates), with values shown as the average of three technical replicates (i.e., three separate wells per donor). The bars indicate the mean value across all donors, and the error bars show the standard deviation.

The changes in the expression of genes identified by RNAseq were also confirmed, with an upregulation of *CXCL8*, *CXCL1*, *IL18* and *COL8A1*, and a downregulation of *CXCL12* by DEX (Fig. 4C).

Cytokine secretion and the effects of hBMSC conditioned medium collected at day 7 were evaluated on hPBMCs. At the protein level, IL-18 concentration in conditioned medium was below the detection limit (data not shown). The trends in IL-8 secretion follows those in *CXCL8* gene expression, with a protein production increased at day 7 in DEX10 compared to AA-2P, and no further increase was observed with DEX100 (Fig. 5).

**Fig. 5.**
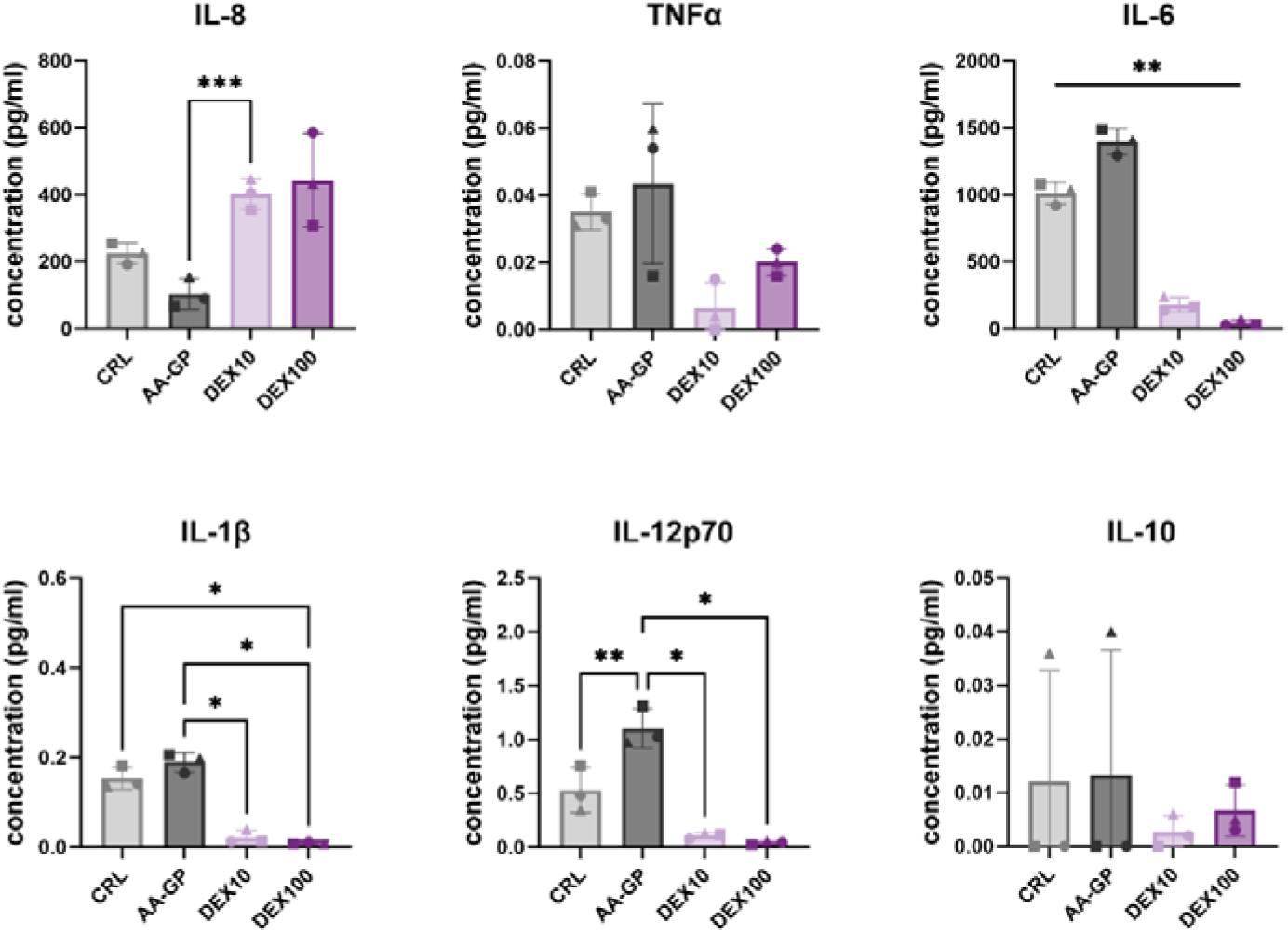
Cytokine production by hBMSCs, 7 days after exposure to DEX. Secretion of cytokines was measured in the conditioned medium of hBMSCs at day 7 by ELISA (IL-8) or multiplex assay (TNFα, IL-6, IL-1β, IL-12p70, IL-10). * p<0.05; ** p<0.01; *** p<0.001. n=3 donors. Each data point represents an individual donor (n = 3 biological replicates), with values shown as the average of two technical replicates (i.e., two separate wells per donor). The bars indicate the mean value across all donors, and the error bars show the standard deviation.

The production of other cytokines was also tested (TNF-α, IL-6, IL-1B, IL-12p70 and IL-10), to evaluate them as a baseline for hPBMC treatment (Fig. 5). hBMSC production of IL-6 decreased with increasing concentration of DEX, correlating with transcript levels. IL-1β and IL-12p70 also followed the same trends though their concentrations were generally very low. The treatment with 10 nM DEX showed a non-significant trend towards a reduction in the concentrations of TNF-α and IL-10; however, the overall levels of these cytokines also remained quite low.

After exposure to hBMSC conditioned medium, hPBMC responses in terms of cytokine production and proliferation were assessed. In both cases, a similar profile was observed for PBMCs treated with MSC conditioned medium or with control medium, indicating no overtly pro- or anti-inflammatory effects of the BMSC secretome (Fig. 6). IL-6 and IL-12p70 levels were mostly determined by their amount secreted by MSCs, therefore they were not taken into consideration for the analysis of PBMC responses. DEX significantly affected hPBMC proliferation, as determined by a two-way ANOVA (*p*_<_0.01), indicating a main effect of treatment across donors.

**Figure 6.**
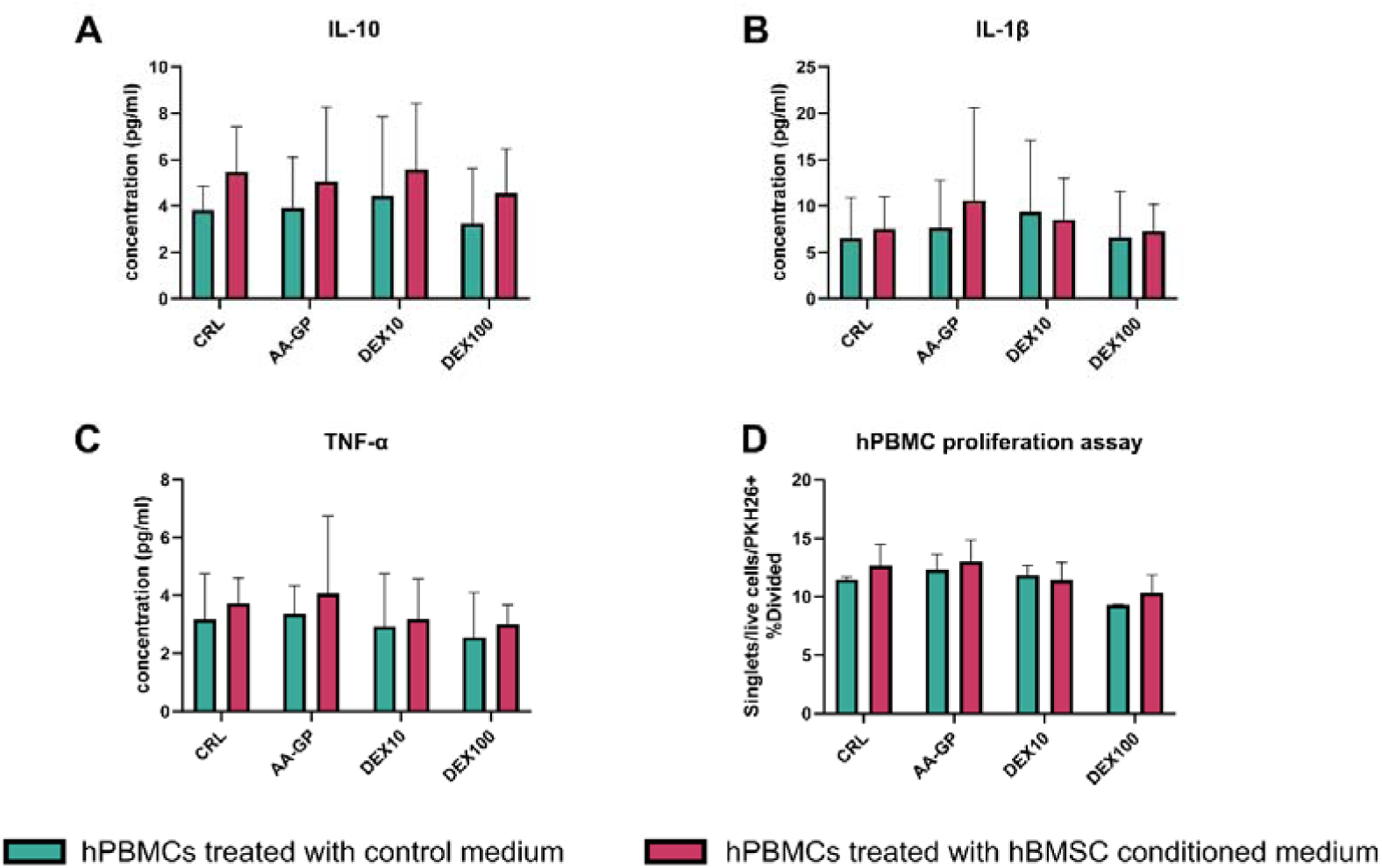
hPBMC responses in terms of **A-C**) cytokine production and **D**) proliferation. The graphs summarize the data from n=3 hBMSC donors and n=3 hPBMC donors, resulting in nine unique donor combinations for PBMCs treated with BMSC conditioned medium. Bars indicate the mean value across all donors, and the error bars show the standard deviation.

## DISCUSSION

This study examined the transcriptomic response of hBMSCs to DEX under pro-osteogenic conditions. By comparing gene expression profiles following DEX and (+)-ZK216348 treatment, we explored potential differences in gene regulation that may involve transactivation or transrepression mechanisms. The overall aim was to identify novel pathways that might explain the inconsistent results often observed in *in vitro* experiments and align them with the clinical effects of glucocorticoids. Although DEX is commonly used to induce osteoblast differentiation at concentration of either 10 or 100 nM [3], emerging evidence suggests that its effects are more intricate and diverse than previously understood. In jaw periosteal cells, for instance, the use of DEX results in lower mineralization and decreased gene expression levels of key osteogenic markers, in comparison to a DEX-free osteogenic medium [22]. The decrease in the osteogenic potential induced by DEX is associated with an increase in the secretion of osteoclastogenic factors and in the expression of adipogenic markers, matching previous observations in BMSCs [22]. Additionally, DEX upregulates *PPARG*, promotes adipogenesis, and inhibits osteocalcin production in BMSCs [12]. This study further explores how DEX influences expression levels of genes involved in several biological processes. Genes related to ossification were both up and downregulated. Some of these regulatory changes align with the promotion of osteogenic differentiation, such as for *ZBTB16* upregulation [23] or for *SFRP4* downregulation [24], while other genes are regulated the opposite way, as in the case for the downregulation of *POSTN* [25,26].

Similar behaviour is observed in genes related to inflammation and to extracellular matrix organization. Along with the expected downregulation of proinflammatory mediators such as IL-6, other pro-inflammatory genes were unexpectedly upregulated by DEX. A notable example of this is the SAA1 gene (controlled by transactivation), which encodes the acute phase protein Serum Amyloid A1. SAA proteins are associated to senescence and bone loss in several conditions [27–30]. The upregulation of *CXCL1*, *CXCL8*, and *IL18* was consistent with a transactivation-associated regulatory pattern, as suggested by comparative analysis with (+)-ZK216348. This was further supported by gene expression data, with *CXCL8* also validated at the protein level, indicating that DEX-treated hBMSCs exhibited enhanced IL-8 secretion. The higher expression and secretion of IL-8 in this scenario does not seem to rely on IL-1β or IL-6, which are recognized for their role in promoting IL-8 expression [31,32]. Instead, these cytokines are significantly downregulated by DEX, thereby indirectly validating the transactivation influence of DEX in regulating *CXCL8* expression. In 1996, Chaudhary and Avioli showed that DEX and other glucocorticoids are able to inhibit IL-1β mediated stimulation of IL-8 production [31]. However, their experiments were not performed in an osteogenic-promoting environment. In contrast, other researchers used a medium containing β-glycerophosphate and ascorbic acid, observing higher IL-8 expression during differentiation in a medium supplemented with 100 nM DEX [33]. This variation in responses highlights the context-dependent effects of DEX and emphasizes the importance of carefully considering experimental differences when comparing results across the literature.

Besides these considerations, it is striking that DEX promotes *CXCL8* expression during osteogenic differentiation of hBMSCs. Though CXCL8/IL-8 is a pleiotropic chemokine, it is primarily involved in immune responses and mainly acts as a pro-inflammatory mediator and a neutrophil chemoattractant [34]. As such, high IL-8 levels are associated with chronic inflammatory disorders, including rheumatoid arthritis [35]. However, its role in progenitor cell differentiation or bone biology remains unclear. Several studies suggest that elevated IL-8 has a negative influence on bone homeostasis. IL-8 expressed during BMSC osteogenic differentiation mediates the recruitment of CD4^+^T cells and inhibits the polarization of Treg cells, leading to a pro-inflammatory environment [36]. IL-8 is also known to promote osteoclast formation [37,38] and a study suggested that the increased expression of IL-8 in jaw periosteal cells in the presence of DEX may contribute to the osteoclastogenic inducing effects observed [22]. There is also a possible autocrine effect on osteoblasts further increasing osteoblast-mediated osteoclastogenesis [39]. Furthermore, IL-8 is produced by adipocytes [40], and in femur marrow adipose tissue it is associated to processes of inflamm-aging [41]. High levels of this chemokine have been associated with vertebral osteoporotic fractures [42], decreased bone mineral density (BMD) in patients with cystic fibrosis [43], and to be involved together with IL-18 in the pathogenesis of bone disease in Cushing’s syndrome, characterized by elevated cortisol levels [44]. Additionally, IL-8 is one component of the senescence-associated secretory phenotype (SASP), and glucocorticoids induce a bone-SASP and accelerated senescence in glucocorticoid-induced osteoporosis [45]. On the other hand, IL-8 might also have beneficial effects on bone healing. Indeed, it has proangiogenic properties, and it can stimulate vascular endothelial growth factor (VEGF) secretion [46]. However, in hBMSCs treated with DEX, VEGF expression is usually reduced [47]. Studies have shown that IL-8 enhances cell recruitment and improves both endochondral bone formation and osteochondral regeneration [48,49]. Thus, CXCL8/IL-8 may support bone healing in contexts requiring localized inflammation, such as fracture repair, but could disrupt bone homeostasis when chronically elevated or inappropriately activated.

Considering that DEX regulates the expression of both pro- and anti-inflammatory cytokines, it is important to understand the overall balance and the effect they have on triggering immune cell responses. Therefore, the effects of the hBMSC conditioned medium on hPBMCs were assessed. Overall, proliferation and cytokine release assays did not indicate a clear pro- or anti-inflammatory effect of the hBMSC secretome, suggesting a balanced cytokine profile or a counteractive influence of DEX on PBMCs, even though appropriate controls were included to discern this effect. A limitation of this model is the use of unsorted hPBMCs rather than selected immune cell populations, such as T cells or macrophages.

DEX influences the expression of various collagen genes. It reduces *COL14A1* levels via transactivation, while transrepression accounts for the downregulation of *COL10A1* and the increase in *COL8A1* and *COL11A1* levels, as for these genes (+)-ZK216348 showed effects comparable to those of DEX. Type XIV collagen is associated with type I collagen fibrils, and it is important for the maintenance of network organization and for the biomechanical properties of tissues [50,51]. *COL11A1* is also important for extracellular matrix organization and mineralization [52,53]. The role of *COL8A1* expression in bone is much less clear. The results of this study suggest that *COL8A1* expression may be suppressed by a DEX-regulated factor acting through transrepression. *COL8A1* encodes for one of the chains of a short-chain, network forming collagen expressed mainly by corneal and vascular endothelial cells. Type VIII collagen encoding genes (*COL8A1* and *COL8A2*) are paralogues of COL10A1 and all of them are vertebrate-specific, with *Col8a1* expressed in mammalian osteoblasts [54,55]. It is believed that the general function of type VIII collagen is to create structures able to withstand compressive forces [56]. Rapidly proliferating cells *in vitro* appear to secrete high levels of type VIII collagen in the medium, but not quiescent cells [56]. In musculoskeletal tissues, both foetal calf cartilage and calvaria periosteum express this collagen type [57]. Both *Col8a1* and *Col8a2* are more expressed in mouse osteoblasts than osteocytes [55]. In bone repair, *Col8a1* was found to be overexpressed in mouse injury-induced fibrogenic cells derived from periosteal cells responding to fracture [58]. Analogously, in mouse calvarial defects, endogenous fibroblast populations form a thin collagen network containing type VIII collagen prior to calvarial bone ossification, which is then replaced by the typical type I collagen-based matrix [59]. In human cells, upregulation of both messenger and circular RNA transcribed from the *COL8A1* gene during DEX-induced osteogenic differentiation was previously described [60]. Altogether, it is likely that type VIII collagen has an important role in bone formation and repair, though further studies are warranted to understand its role in bone homeostasis and disease and in cell differentiation. These findings highlight the complex and context-dependent regulation of collagen genes by DEX, with potential implications for matrix remodelling during osteogenesis.

## CONCLUSIONS

Overall, the results of this study indicate that DEX modulates several pathways related to ossification, extracellular matrix organization, and inflammation. This study reveals that, although the anti-inflammatory properties of DEX are typically attributed to transrepression, in hBMSCs DEX also enhances the expression of certain pro-inflammatory mediators. While it is suggested that transactivation mechanisms may be involved, further studies are needed to confirm the underlying regulatory pathways using direct mechanistic assays. The role of key factors regulated by DEX, such as IL-8, IL-18, and type VIII collagen, is currently not clear and need further investigation. It is essential to determine whether these factors are part of the physiological processes of either bone formation or homeostasis, or if they represent side effects of excessive use of glucocorticoids. If this is the case, this study can clarify some of the detrimental effects of excess glucocorticoids on hBMSCs, providing useful insights on the pathophysiology of glucocorticoid-induced osteoporosis. Furthermore, even though our model did not show a pro- or anti-inflammatory effect of the hBMSC conditioned medium, the pattern of cytokine expression suggests that the interaction of osteoprogenitors with immune cells can play an important role in determining the effects of glucocorticoids on bone, especially at a local level.

In conclusion, the application of DEX for promoting osteogenic differentiation *in vitro* warrants re-evaluation. Embracing more physiologically relevant methods could significantly enhance translational potential, influencing all *in vitro* studies in bone research. Future studies should explore alternative osteogenic induction strategies that better mimic physiological conditions and minimize off-target inflammatory effects.

## DECLARATIONS

### Ethics approval and consent to participate

Bone marrow was collected from patients undergoing spine surgery at the Inselspital Bern with signed informed patient consent. The Swiss Human Research Act does not apply to research that utilizes anonymized biological material and/or anonymously collected or anonymized health-related data. Therefore, this project did not need to be approved by an ethics committee. Patients’ general consent was obtained, which also covers the anonymization of health-related data and biological material. The study was conducted in accordance with the Declaration of Helsinki.

### Consent for publication

Not applicable.

### Availability of data and materials

Sequencing data are available online at: http://www.ncbi.nlm.nih.gov/bioproject/1244756. Other data are available from the corresponding author on reasonable request.

### Authors’ contributions

**ABD**: Methodology, Formal analysis, Investigation, Data Curation, Writing - Review & Editing, Visualization, Funding acquisition. **CS**: Methodology, Validation, Investigation, Writing - Review & Editing. **JUG**: Validation, Writing - Review & Editing. **CL**: Investigation. **EP**: Investigation. **GG**: Methodology, Funding acquisition. **SH**: Resources, Writing - Review & Editing. **MJS**: Conceptualization, Funding acquisition, Writing - Review & Editing. **EDB**: Conceptualization, Methodology, Formal analysis, Data Curation, Writing - Original Draft, Visualization, Supervision, Project administration, Funding acquisition. All the authors approved the final version of the manuscript.

## Acknowledgements

Figure 1 was created with BioRender.com.

