## Supplementary file 1 for "Modulation of ossification and inflammatory pathways during dexamethasone-induced *in vitro* osteogenesis"

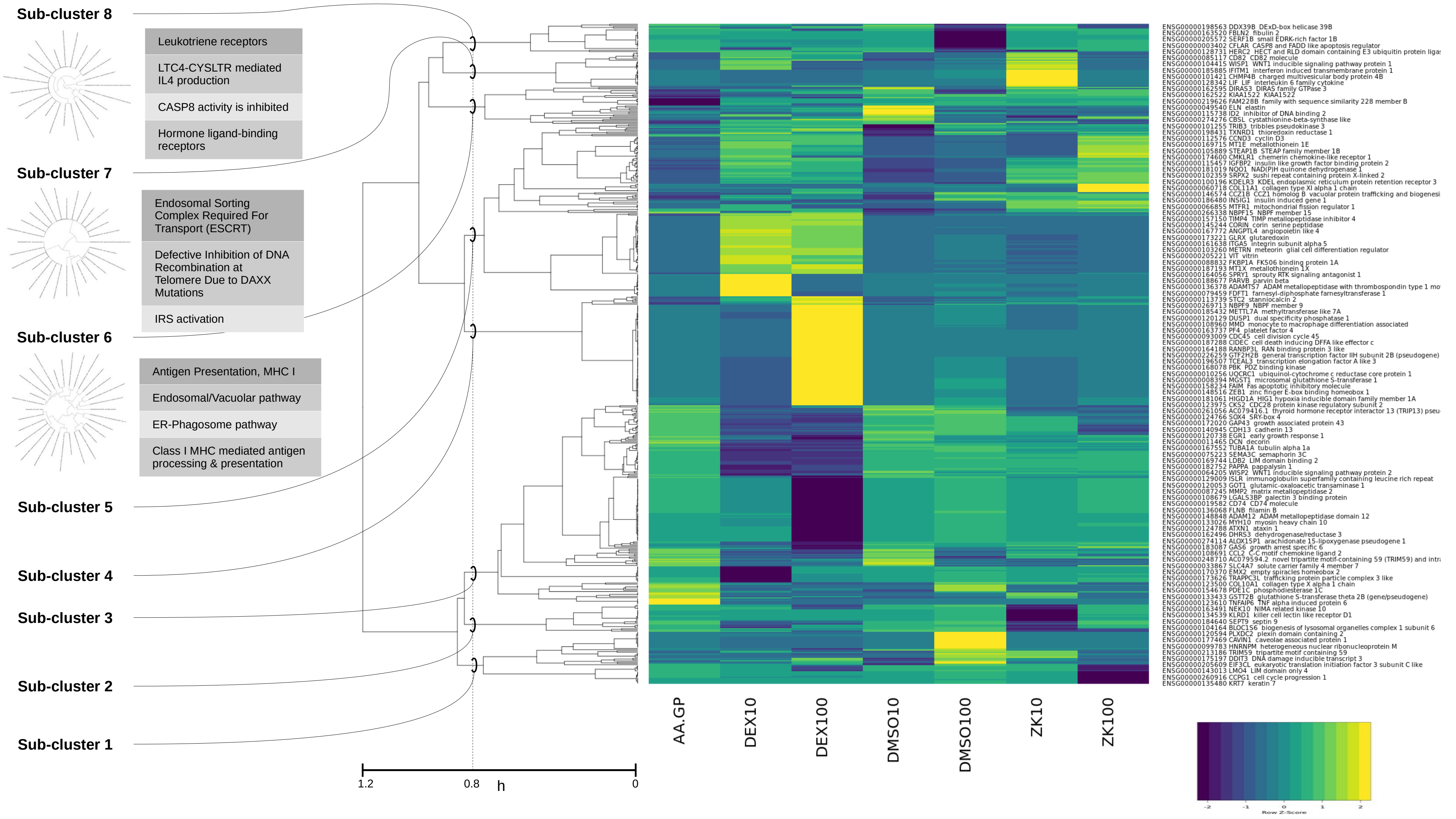
Sub-cluster 8
Leukotriene receptors
LTC4-CYSLTR mediated IL4 production
CASP8 activity is inhibited
Hormone ligand-binding receptors
Sub-cluster 7
Endosomal Sorting Complex Required For Transport (ESCRT)
Defective Inhibition of DNA Recombination at Telomere Due to DAXX Mutations
IRS activation
Sub-cluster 6
Antigen Presentation, MHC I
Endosomal/Vacuolar pathway
ER-Phagosome pathway
Class I MHC mediated antigen processing & presentation
Sub-cluster 5
Sub-cluster 4
Sub-cluster 3
Sub-cluster 2
Sub-cluster 1
1.2
0.8
h
0
AA.GP
DEX10
DEX100
DMSO10
DMSO100
ZK10
ZK100
ENSG00000198563 DDX39B DExD-box helicase 39B
ENSG00000163520 FBLN2 fibulin 2
ENSG00000205572 SERF1B small EDRK-rich factor 1B
ENSG00000003402 CFLAR CASP8 and FADD like apoptosis regulator
ENSG00000128731 HERC2 HECT and RLD domain containing E3 ubiquitin protein ligas
ENSG00000085117 CD82 CD82 molecule
ENSG00000104415 WISP1 WNT1 inducible signaling pathway protein 1
ENSG00000185885 IFITM1 interferon induced transmembrane protein 1
ENSG00000101421 CHMP4B charged multivesicular body protein 4B
ENSG00000128342 LIF LIF interleukin 6 family cytokine
ENSG00000162595 DIRAS3 DIRAS family GTPase 3
ENSG00000162522 KIAA1522 KIAA1522
ENSG00000219626 FAM228B family with sequence similarity 228 member B
ENSG00000049540 ELN elastin
ENSG00000115738 ID2 inhibitor of DNA binding 2
ENSG00000274276 CBSL cystathionine-beta-synthase like
ENSG00000101255 TRIB3 tribbles pseudokinase 3
ENSG00000198431 TXNRD1 thioredoxin reductase 1
ENSG00000112576 CCND3 cyclin D3
ENSG00000169715 MT1E metallothionein 1E
ENSG00000105889 STEAP1B STEAP family member 1B
ENSG00000174600 CMKLR1 chemerin chemokine-like receptor 1
ENSG00000115457 IGFBP2 insulin like growth factor binding protein 2
ENSG00000181019 NQO1 NAD(P)H quinone dehydrogenase 1
ENSG00000102359 SRPX2 sushi repeat containing protein X-linked 2
ENSG00000100196 KDELR3 KDEL endoplasmic reticulum protein retention receptor 3
ENSG00000060718 COL11A1 collagen type XI alpha 1 chain
ENSG00000146574 CCZ1B CCZ1 homolog B vacuolar protein trafficking and biogenesi
ENSG00000186480 INSIG1 insulin induced gene 1
ENSG00000066855 MTFR1 mitochondrial fission regulator 1
ENSG00000266338 NBPF15 NBPF member 15
ENSG00000157150 TIMP4 TIMP metallopeptidase inhibitor 4
ENSG00000145244 CORIN corin serine peptidase
ENSG00000167772 ANGPTL4 angiopoietin like 4
ENSG00000173221 GLRX glutaredoxin
ENSG00000161638 ITGA5 integrin subunit alpha 5
ENSG00000103260 METRN meteorin glial cell differentiation regulator
ENSG00000205221 VIT vitrin
ENSG00000088832 FKBP1A FK506 binding protein 1A
ENSG00000187193 MT1X metallothionein 1X
ENSG00000164056 SPRY1 sprouty RTK signaling antagonist 1
ENSG00000188677 PARVB parvin beta
ENSG00000136378 ADAMTS7 ADAM metallopeptidase with thrombospondin type 1 mo
ENSG00000079459 FDFT1 farnesyl-diphosphate farnesyltransferase 1
ENSG00000113739 STC2 stanniocalcin 2
ENSG00000269713 NBPF9 NBPF member 9
ENSG00000185432 METTL7A methyltransferase like 7A
ENSG00000120129 DUSP1 dual specificity phosphatase 1
ENSG00000108960 MMD monocyte to macrophage differentiation associated
ENSG00000163737 PF4 platelet factor 4
ENSG00000093009 CDC45 cell division cycle 45
ENSG00000187288 CIDEC cell death inducing DFFA like effector c
ENSG00000164188 RANBP3L RAN binding protein 3 like
ENSG00000226259 GTF2H2B general transcription factor IIH subunit 2B (pseudogene)
ENSG00000196507 TCEAL3 transcription elongation factor A like 3
ENSG00000168078 PBK PDZ binding kinase
ENSG00000010256 UQCRC1 ubiquinol-cytochrome c reductase core protein 1
ENSG00000008394 MGST1 microsomal glutathione S-transferase 1
ENSG00000158234 FAIM Fas apoptotic inhibitory molecule
ENSG00000148516 ZEB1 zinc finger E-box binding homeobox 1
ENSG00000181061 HIGD1A HIG1 hypoxia inducible domain family member 1A
ENSG00000123975 CKS2 CDC28 protein kinase regulatory subunit 2
ENSG00000261056 AC079416.1 thyroid hormone receptor interactor 13 (TRIP13) pseu
ENSG00000124766 SOX4 SRY-box 4
ENSG00000172020 GAP43 growth associated protein 43
ENSG00000140945 CDH13 cadherin 13
ENSG00000120738 EGR1 early growth response 1
ENSG00000011465 DCN decorin
ENSG00000167552 TUBA1A tubulin alpha 1a
ENSG00000075223 SEMA3C semaphorin 3C
ENSG00000169744 LDB2 LIM domain binding 2
ENSG00000182752 PAPPA pappalysin 1
ENSG00000064205 WISP2 WNT1 inducible signaling pathway protein 2
ENSG00000129009 ISLR immunoglobulin superfamily containing leucine rich repeat
ENSG00000120053 GOT1 glutamic-oxaloacetic transaminase 1
ENSG00000087245 MMP2 matrix metallopeptidase 2
ENSG00000108679 LGALS3BP galectin 3 binding protein
ENSG00000019582 CD74 CD74 molecule
ENSG00000136068 FLNB filamin B
ENSG00000148848 ADAM12 ADAM metallopeptidase domain 12
ENSG00000133026 MYH10 myosin heavy chain 10
ENSG00000124788 ATXN1 ataxin 1
ENSG00000162496 DHRS3 dehydrogenase/reductase 3
ENSG00000274114 ALOX15P1 arachidonate 15-lipoxygenase pseudogene 1
ENSG00000183087 GAS6 growth arrest specific 6
ENSG00000108691 CCL2 C-C motif chemokine ligand 2
ENSG00000248710 AC079594.2 novel tripartite motif-containing 59 (TRIM59) and intra
ENSG00000033867 SLC4A7 solute carrier family 4 member 7
ENSG00000170370 EMX2 empty spiracles homeobox 2
ENSG00000173626 TRAPPC3L trafficking protein particle complex 3 like
ENSG00000123500 COL10A1 collagen type X alpha 1 chain
ENSG00000154678 PDE1C phosphodiesterase 1C
ENSG00000133433 GSTT2B glutathione S-transferase theta 2B (gene/pseudogene)
ENSG00000123610 TNFAIP6 TNF alpha induced protein 6
ENSG00000163491 NEK10 NIMA related kinase 10
ENSG00000134539 KLRD1 killer cell lectin like receptor D1
ENSG00000184640 SEPT9 septin 9
ENSG00000104164 BLOC1S6 biogenesis of lysosomal organelles complex 1 subunit 6
ENSG00000120594 PLXDC2 plexin domain containing 2
ENSG00000177469 CAVIN1 caveolae associated protein 1
ENSG00000099783 HNRNPM heterogeneous nuclear ribonucleoprotein M
ENSG00000213186 TRIM59 tripartite motif containing 59
ENSG00000175197 DDIT3 DNA damage inducible transcript 3
ENSG00000205609 EIF3CL eukaryotic translation initiation factor 3 subunit C like
ENSG00000143013 LMO4 LIM domain only 4
ENSG00000260916 CCPG1 cell cycle progression 1
ENSG00000135480 KRT7 keratin 7
-2
-1
0
1
2
Row Z-Score

### Sub-cluster 1

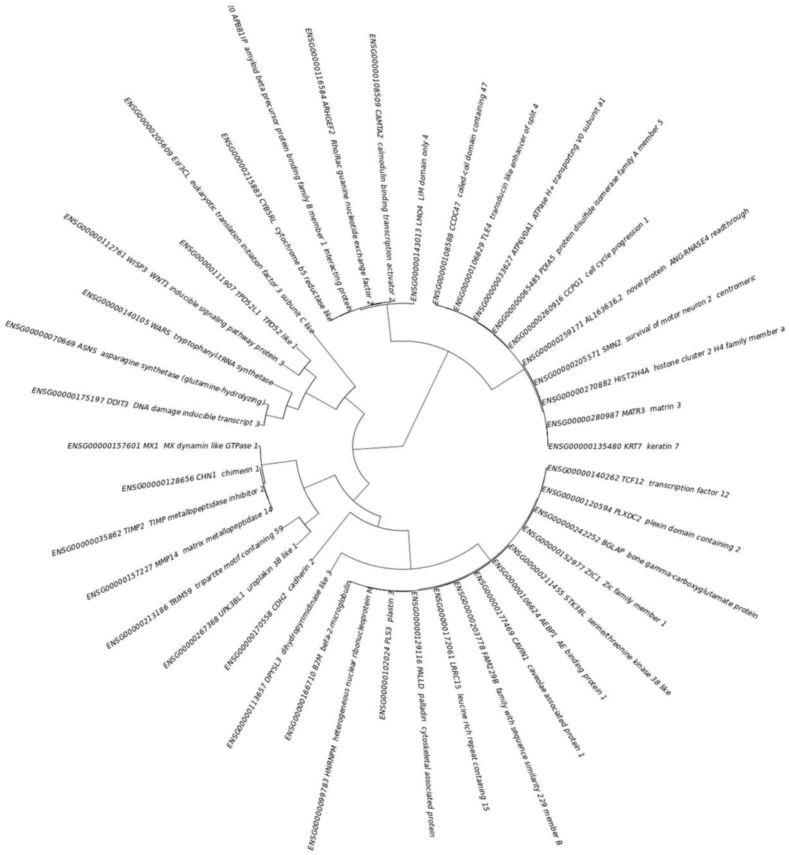

### 25 most relevant pathways sorted by p-value (https://reactome.org/)

| Pathway name | Submitted entities found |
|---|---|
| Response of EIF2AK1 (HRI) to heme deficiency | DDIT3;ASNS |
| ATF4 activates genes in response to endoplasmic reticulum stress | DDIT3;ASNS |
| PERK regulates gene expression | DDIT3;ASNS |
| Response of EIF2AK4 (GCN2) to amino acid deficiency | DDIT3;ASNS |
| Cellular response to starvation | DDIT3;ASNS |
| Activation of Matrix Metalloproteinases | MMP14;TIMP2 |
| RHOB GTPase cycle | CAVIN1;ARHGEF2 |
| Maturation of protein M | |
| Virion Assembly and Release | |
| Myogenesis | CDH2;TCF12 |
| Regulation of gene expression in endocrine-committed (NEUROG3+) progenitor cells | |
| GLI proteins bind promoters of Hh responsive genes to promote transcription | |
| RHOA GTPase cycle | CAVIN1;ARHGEF2 |
| Transport of gamma-carboxylated protein precursors from the endoplasmic reticulum to the Golgi apparatus | BGLAP |
| Nef mediated downregulation of MHC class I complex cell surface expression | B2M |
| TRAF6 mediated IRF7 activation in TLR7/8 or 9 signaling | |
| Erythrocytes take up carbon dioxide and release oxygen | CYB5RL |
| O2/CO2 exchange in erythrocytes | CYB5RL |
| Transcription of SARS-CoV-1 sgRNAs | |
| Regulation of BACH1 activity | |
| Insulin receptor recycling | ATP6V0A1 |
| Maturation of protein 3a | |
| Gamma-carboxylation of protein precursors | BGLAP |
| Reduction of cytosolic Ca++ levels | |
| Removal of aminoterminal propeptides from gamma-carboxylated proteins | BGLAP |

### Sub-cluster 2

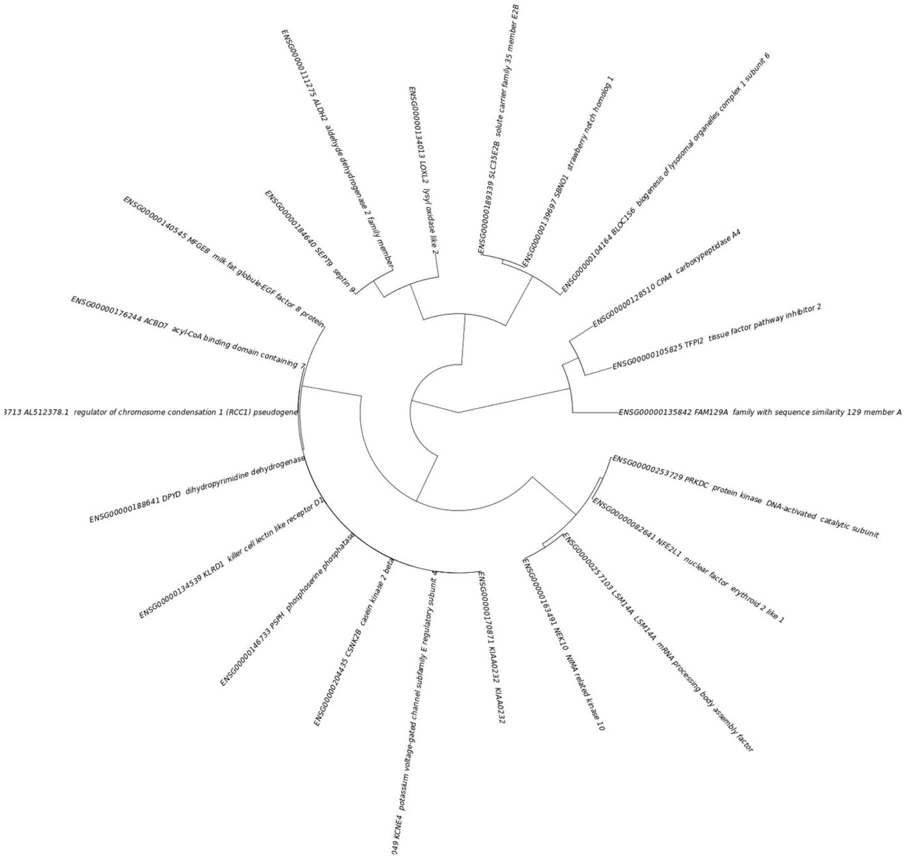


### 25 most relevant pathways sorted by p-value (https://reactome.org/)

| Pathway name | Submitted entities found |
|---|---|
| Crosslinking of collagen fibrils | LOXL2 |
| Elastic fibre formation | LOXL2 |
| Assembly of collagen fibrils and other multimeric structures | LOXL2 |
| Phase 3 - rapid repolarisation | KCNE4 |
| Metabolism of serotonin | ALDH2 |
| Condensation of Prometaphase Chromosomes | CSNK2B |
| Serotonin clearance from the synaptic cleft | ALDH2 |
| Collagen formation | LOXL2 |
| Phase 2 - plateau phase | KCNE4 |
| Nef mediated downregulation of MHC class I complex cell surface expression | |
| Serine biosynthesis | PSPH |
| Ethanol oxidation | ALDH2 |
| FBXW7 Mutants and NOTCH1 in Cancer | |
| Loss of Function of FBXW7 in Cancer and NOTCH1 Signaling | |
| Constitutive Signaling by NOTCH1 t(7;9)(NOTCH1:M1580_K2555) Translocation Mutant | |
| Signaling by NOTCH1 t(7;9)(NOTCH1:M1580_K2555) Translocation Mutant | |
| Signaling by NOTCH1 HD Domain Mutants in Cancer | |
| Constitutive Signaling by NOTCH1 HD Domain Mutants | |
| Fibronectin matrix formation | |
| Cooperation of PDCL (PhLP1) and TRiC/CCT in G-protein beta folding | CSNK2B |
| Receptor Mediated Mitophagy | CSNK2B |
| Pyrimidine catabolism | DPYD |
| PIPs transport between late endosome and Golgi membranes | |
| PIPs transport between early endosome and Golgi membranes | |
| PIPs transport between early and late endosome membranes | |

### Sub-cluster 3

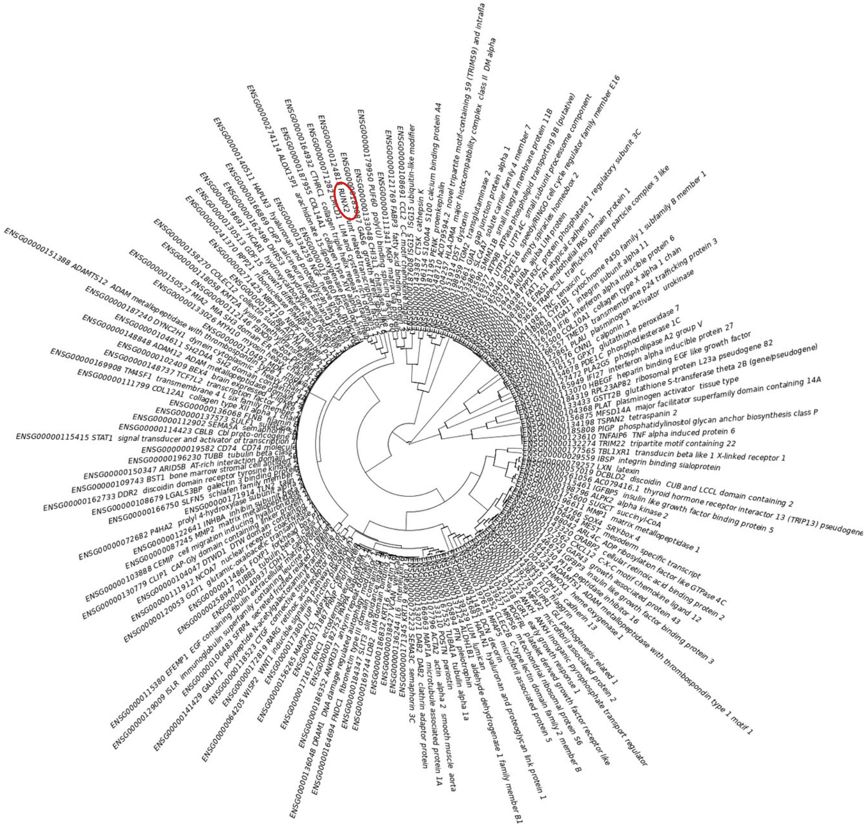

### 25 most relevant pathways sorted by p-value (https://reactome.org/)

| Pathway name | Submitted entities found |
|---|---|
| Collagen degradation | COL14A1;MMP1;CTSK;MMP2;COL12A1;COL10A1;PLAT |
| Degradation of the extracellular matrix | ADAMTS1;COL14A1;MMP1;CTSK;MMP2;COL12A1;COL10A1;PLAT;DCN |
| Activation of Matrix Metalloproteinases | MMP1;CTSK;MMP2;PLAT |
| Microtubule-dependent trafficking of connexons from Golgi to the plasma membrane | GJA1;TUBA1A;TUBB3;TUBB |
| Transport of connexons to the plasma membrane | GJA1;TUBA1A;TUBB3;TUBB |
| Interleukin-4 and Interleukin-13 signaling | IL6;MMP1;STAT1;MMP2;CCL2;HMOX1 |
| Post-translational protein phosphorylation | IL6;VCAN;IGFBP5;IGFBP3;PENK;TNC;GAS6 |
| Extracellular matrix organization | DST;COL14A1;MMP1;LUM;MMP2;COL12A1;TNC;PLAT;DCN;HAPLN1;MFAP5;VCAN;EFEMP1;IBSP;P4HA2;ADAMTS1;CTSK;ITGA11;ADAM12;MFAP2;COL10A1;DDR2 |
| Regulation of Insulin-like Growth Factor (IGF) transport and uptake by Insulin-like Growth Factor Binding Proteins (IGFBPs) | IL6;VCAN;IGFBP5;MMP1;IGFBP3;PAPPA;MMP2;PENK;TNC;GAS6 |
| Signaling by FGFR1 amplification mutants | FLG |
| RHO GTPases activate IQGAPs | CLIP1;TUBA1A;TUBB3;TUBB |
| Assembly of collagen fibrils and other multimeric structures | DST;COL14A1;MMP1;COL12A1;COL10A1 |
| ECM proteoglycans | VCAN;IBSP;LUM;TNC;DCN;HAPLN1 |
| Prefoldin mediated transfer of substrate to CCT/TriC | TUBA1A;TUBB3;TUBB |
| Defective CHST3 causes SEDCJD | VCAN;DCN |
| Defective CHST14 causes EDS, musculocontractural type | VCAN;DCN |
| Formation of tubulin folding intermediates by CCT/TriC | TUBA1A;TUBB3;TUBB |
| Defective CHSY1 causes TPBS | VCAN;DCN |
| Dermatan sulfate biosynthesis | VCAN;DCN |
| Defective B3GALTL causes PpS | SEMA5A;ADAMTS1;ADAMTS12 |
| Post-chaperonin tubulin folding pathway | TUBA1A;TUBB3;TUBB |
| Molecules associated with elastic fibres | MFAP5;EFEMP1;MFAP2 |
| O-glycosylation of TSR domain-containing proteins | SEMA5A;ADAMTS1;ADAMTS12 |
| Collagen chain trimerization | COL14A1;COL12A1;COL10A1 |
| Interferon alpha/beta signaling | EGR1;IFI27;STAT1;IFI6;ISG15 |

### Sub-cluster 4

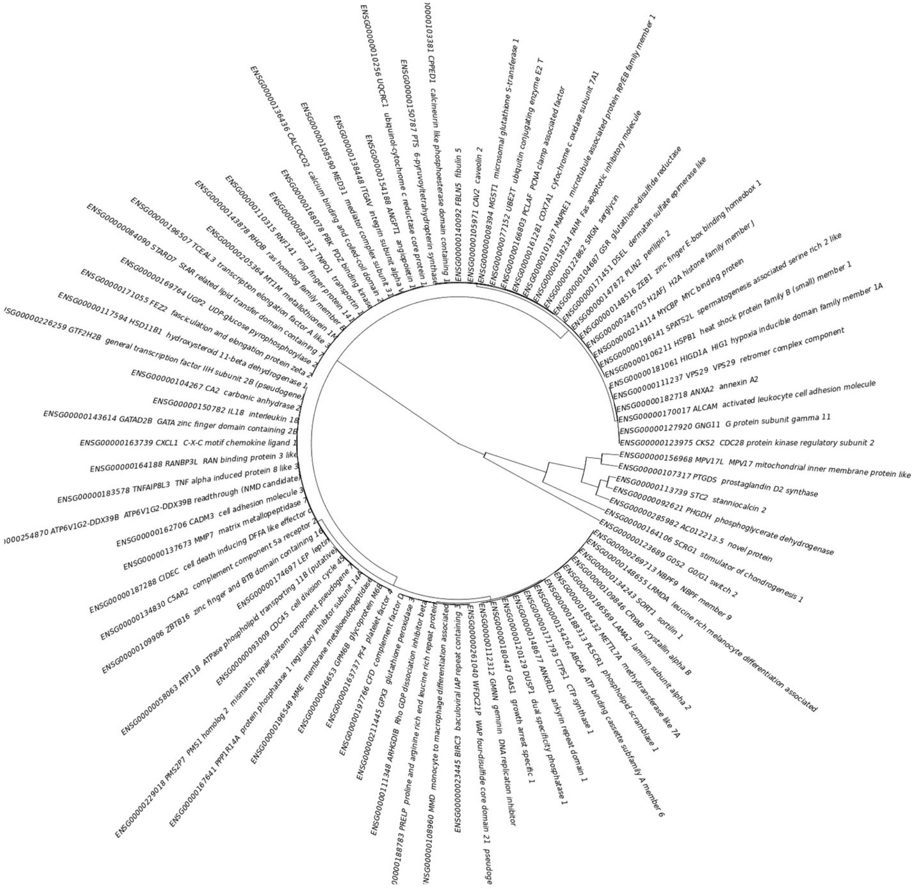

### 25 most relevant pathways sorted by p-value (https://reactome.org/)

| Pathway name | Submitted entities found |
|---|---|
| Interleukin-18 signaling | IL18 |
| Reversible hydration of carbon dioxide | CA2 |
| Neutrophil degranulation | CFD;MME;ANXA2;MMD;MGST1;CXCL1;ATP11B;ITGAV;METTL7A;CPPED1 |
| Molecules associated with elastic fibres | ITGAV;FBLN5 |
| Elastic fibre formation | ITGAV;FBLN5 |
| Erythrocytes take up oxygen and release carbon dioxide | CA2 |
| TNF receptor superfamily (TNFSF) members mediating non-canonical NF-kB pathway | BIRC3 |
| O2/CO2 exchange in erythrocytes | CA2 |
| Erythrocytes take up carbon dioxide and release oxygen | CA2 |
| Laminin interactions | LAMA2;ITGAV |
| TICAM1, RIP1-mediated IKK complex recruitment | BIRC3 |
| Interleukin-10 signaling | IL18;CXCL1 |
| Chemokine receptors bind chemokines | CXCL1;PF4 |
| IKK complex recruitment mediated by RIP1 | BIRC3 |
| NTF3 activates NTRK3 signaling | |
| Metallothioneins bind metals | MT1M |
| Interconversion of nucleotide di- and triphosphates | GSR;CTPS1 |
| Defective B4GALT1 causes B4GALT1-CDG (CDG-2d) | PRELP |
| Defective ST3GAL3 causes MCT12 and EIEE15 | PRELP |
| Defective CHST6 causes MCDC1 | PRELP |
| HSF1-dependent transactivation | HSPB1;CRYAB |
| Transcriptional regulation of testis differentiation | PTGDS |
| Activation of the pre-replicative complex | CDC45;GMNN |
| NGF processing | |
| Expression and Processing of Neurotrophins | |

### Sub-cluster 5

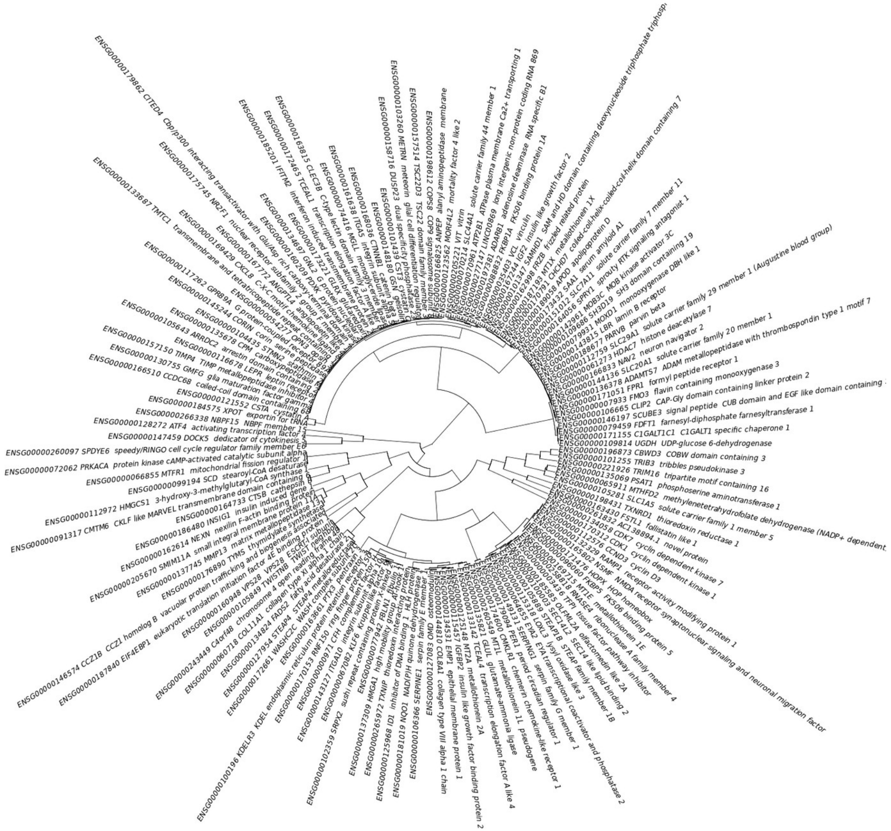

### 25 most relevant pathways sorted by p-value (https://reactome.org/)

| Pathway name | Submitted entities found |
| --- | --- |
| RUNX2 regulates genes involved in cell migration | MMP13;ITGA5 |
| Collagen degradation | MMP13;COL11A1;COL8A1;CTSB |
| Assembly of collagen fibrils and other multimeric structures | MMP13;COL11A1;LOXL3;COL8A1;CTSB |
| Formyl peptide receptors bind formyl peptides and many other ligands | FPR1;SAA1 |
| PKA-mediated phosphorylation of key metabolic factors | PRKACA |
| Regulation of cholesterol biosynthesis by SREBP (SREBF) | HMGCS1;SCD;INSIG1;LBR;FDFT1 |
| Activation of gene expression by SREBF (SREBP) | HMGCS1;SCD;LBR;FDFT1 |
| Neutrophil degranulation | CST3;PDXK;CXCL8;GSN;GMFG;ANPEP;CMTM6;FPR1;PTX3;VCL;CTSB |
| Degradation of the extracellular matrix | SCUBE3;MMP13;COL11A1;COL8A1;CTSB |
| Defective binding of RB1 mutants to E2F1,(E2F2, E2F3) | CCND3;CDK1 |
| Aberrant regulation of mitotic G1/S transition in cancer due to RB1 defects | CCND3;CDK1 |
| Platelet degranulation | CLEC3B;CXCL8;SERPINE1;IGF2;SERPING1;VCL |
| Regulation of glycolysis by fructose 2,6-bisphosphate metabolism | PRKACA |
| Response to elevated platelet cytosolic Ca2+ | CLEC3B;CXCL8;SERPINE1;IGF2;SERPING1;PRKACA;VCL |
| Metallothioneins bind metals | MT2A;MT1X;MT1E |
| Collagen formation | MMP13;COL11A1;LOXL3;COL8A1;CTSB |
| Insulin-like Growth Factor-2 mRNA Binding Proteins (IGF2BPs/IMPs/VICKZs) bind RNA | IGF2 |
| Elastic fibre formation | LOXL3;FBLN1;ITGA5 |
| Defective SERPING1 causes hereditary angioedema | SERPING1 |
| Response to metal ions | MT2A;MT1X;MT1E |
| PPARA activates gene expression | HMGCS1;TXNRD1;TRIB3;ANGPTL4;FDFT1 |
| TGFBR1 LBD Mutants in Cancer | FKBP1A;FPR1 |
| C6 deamination of adenosine | ADARB1 |
| Depolymerisation of the Nuclear Lamina | CDK1;PRKACA |
| Phosphorylation of proteins involved in the G2/M transition by Cyclin A:Cdc2 complexes | CDK1 |

### Sub-cluster 6

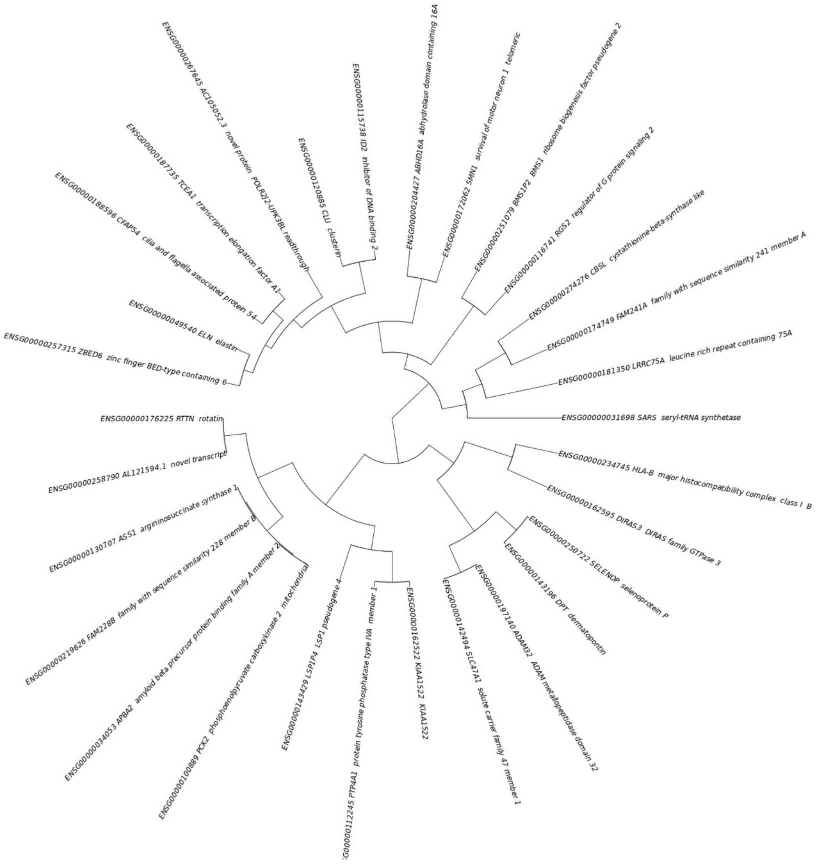


### 25 most relevant pathways sorted by p-value (https://reactome.org/)

| Pathway name | Submitted entities found |
|---|---|
| Antigen Presentation: Folding, assembly and peptide loading of class I MHC | HLA-B |
| Endosomal/Vacuolar pathway | HLA-B |
| ER-Phagosome pathway | HLA-B |
| Class I MHC mediated antigen processing & presentation | HLA-B |
| Antigen processing-Cross presentation | HLA-B |
| Interferon gamma signaling | HLA-B |
| Interferon alpha/beta signaling | HLA-B |
| Immunoregulatory interactions between a Lymphoid and a non-Lymphoid cell | HLA-B |
| Interferon Signaling | HLA-B |
| Adaptive Immune System | HLA-B |
| Cytokine Signaling in Immune system | HLA-B |
| Immune System | HLA-B;CLU |
| Terminal pathway of complement | CLU |
| Cysteine formation from homocysteine | CBSL |
| Aromatic amines can be N-hydroxylated or N-dealkylated by CYP1A2 | |
| NGF-independant TRKA activation | |
| Urea cycle | ASS1 |
| Activation of TRKA receptors | |
| Modulation by Mtb of host immune system | |
| Tat-mediated HIV elongation arrest and recovery | TCEA1 |
| Pausing and recovery of Tat-mediated HIV elongation | TCEA1 |
| HIV elongation arrest and recovery | TCEA1 |
| Pausing and recovery of HIV elongation | TCEA1 |
| Molecules associated with elastic fibres | ELN |
| Platelet degranulation | SELENOP;CLU |

### Sub-cluster 7

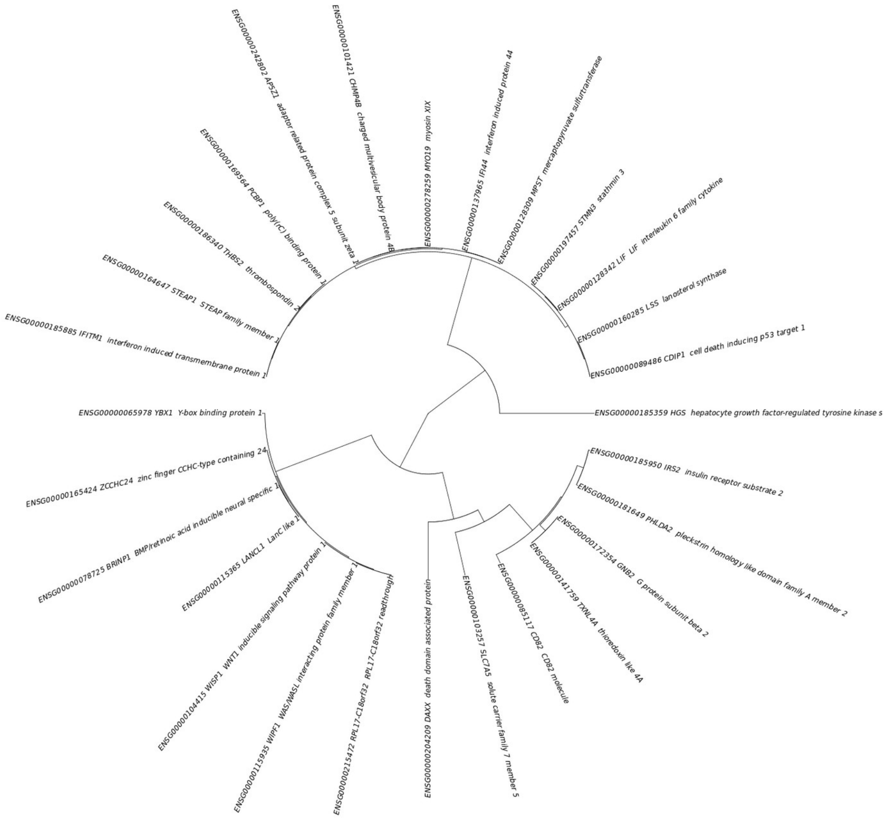


### 25 most relevant pathways sorted by p-value (https://reactome.org/)

| Pathway name | Submitted entities found |
|---|---|
| Endosomal Sorting Complex Required For Transport (ESCRT) | HGS;CHMP4B |
| Defective Inhibition of DNA Recombination at Telomere Due to DAXX Mutations | DAXX |
| IRS activation | IRS2 |
| TFAP2 (AP-2) family regulates transcription of other transcription factors | |
| Erythropoietin activates STAT5 | IRS2 |
| SOS-mediated signalling | IRS2 |
| Evasion of Oxidative Stress Induced Senescence Due to p14ARF Defects | |
| Evasion of Oncogene Induced Senescence Due to p14ARF Defects | |
| Inhibition of Host mRNA Processing and RNA Silencing | |
| RHOT1 GTPase cycle | MYO19 |
| mRNA Splicing - Minor Pathway | YBX1;TXNL4A |
| RHOT2 GTPase cycle | MYO19 |
| Miro GTPase Cycle | MYO19 |
| Insulin-like Growth Factor-2 mRNA Binding Proteins (IGF2BPs/IMPs/VICKZs) bind RNA | |
| Presynaptic function of Kainate receptors | GNB2 |
| Processing of Intronless Pre-mRNAs | |
| Interleukin-10 signaling | LIF |
| G-protein activation | GNB2 |
| ADP signalling through P2Y purinoceptor 12 | GNB2 |
| RHOBTB3 ATPase cycle | HGS |
| Prostacyclin signalling through prostacyclin receptor | GNB2 |
| Activation of gene expression by SREBF (SREBP) | LSS |
| G protein gated Potassium channels | GNB2 |
| Activation of G protein gated Potassium channels | GNB2 |
| Inhibition of voltage gated Ca2+ channels via Gbeta/gamma subunits | GNB2 |

### Sub-cluster 8

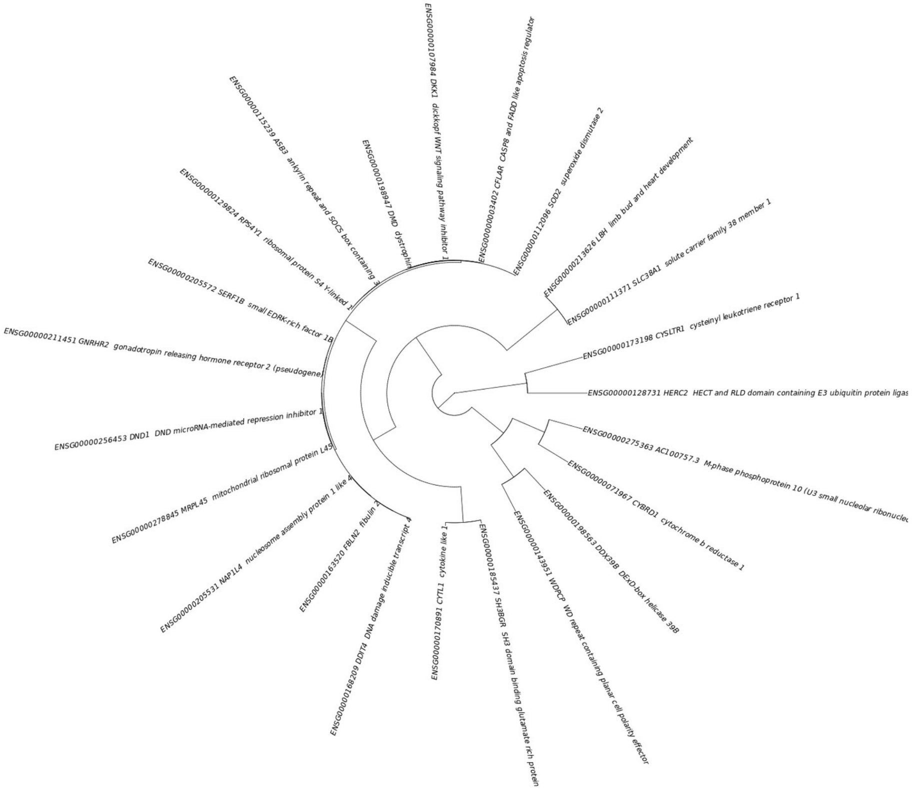


### 25 most relevant pathways sorted by p-value (https://reactome.org/)

| Pathway name | Submitted entities found |
|---|---|
| Leukotriene receptors | CYSLTR1 |
| LTC4-CYSLTR mediated IL4 production | CYSLTR1 |
| CASP8 activity is inhibited | CFLAR |
| Hormone ligand-binding receptors | GNRHR2 |
| Caspase activation via Death Receptors in the presence of ligand | CFLAR |
| Eicosanoid ligand-binding receptors | CYSLTR1 |
| Signaling by LRP5 mutants | DKK1 |
| G alpha (q) signalling events | CYSLTR1;GNRHR2 |
| POU5F1 (OCT4), SOX2, NANOG repress genes related to differentiation | DKK1 |
| RHOBTB2 GTPase cycle | DDX39B |
| Caspase activation via extrinsic apoptotic signalling pathway | CFLAR |
| Negative regulation of TCF-dependent signaling by WNT ligand antagonists | DKK1 |
| TP53 Regulates Metabolic Genes | DDIT4 |
| Class A/1 (Rhodopsin-like receptors) | CYSLTR1;GNRHR2 |
| Dimerization of procaspase-8 | CFLAR |
| Regulation by c-FLIP | CFLAR |
| FOXO-mediated transcription of oxidative stress, metabolic and neuronal genes | SOD2 |
| Neurotransmitter uptake and metabolism In glial cells | SLC38A1 |
| Astrocytic Glutamate-Glutamine Uptake And Metabolism | SLC38A1 |
| Molecules associated with elastic fibres | FBLN2 |
| RHOBTB GTPase Cycle | DDX39B |
| Striated Muscle Contraction | DMD |
| Glucagon-type ligand receptors | CYSLTR1 |
| RNA Polymerase I Transcription Termination | |
| Interactions of Tat with host cellular proteins | |
