## Supplementary figures and images for "Modulation of ossification and inflammatory pathways during dexamethasone-induced *in vitro* osteogenesis"

### Fig. S2

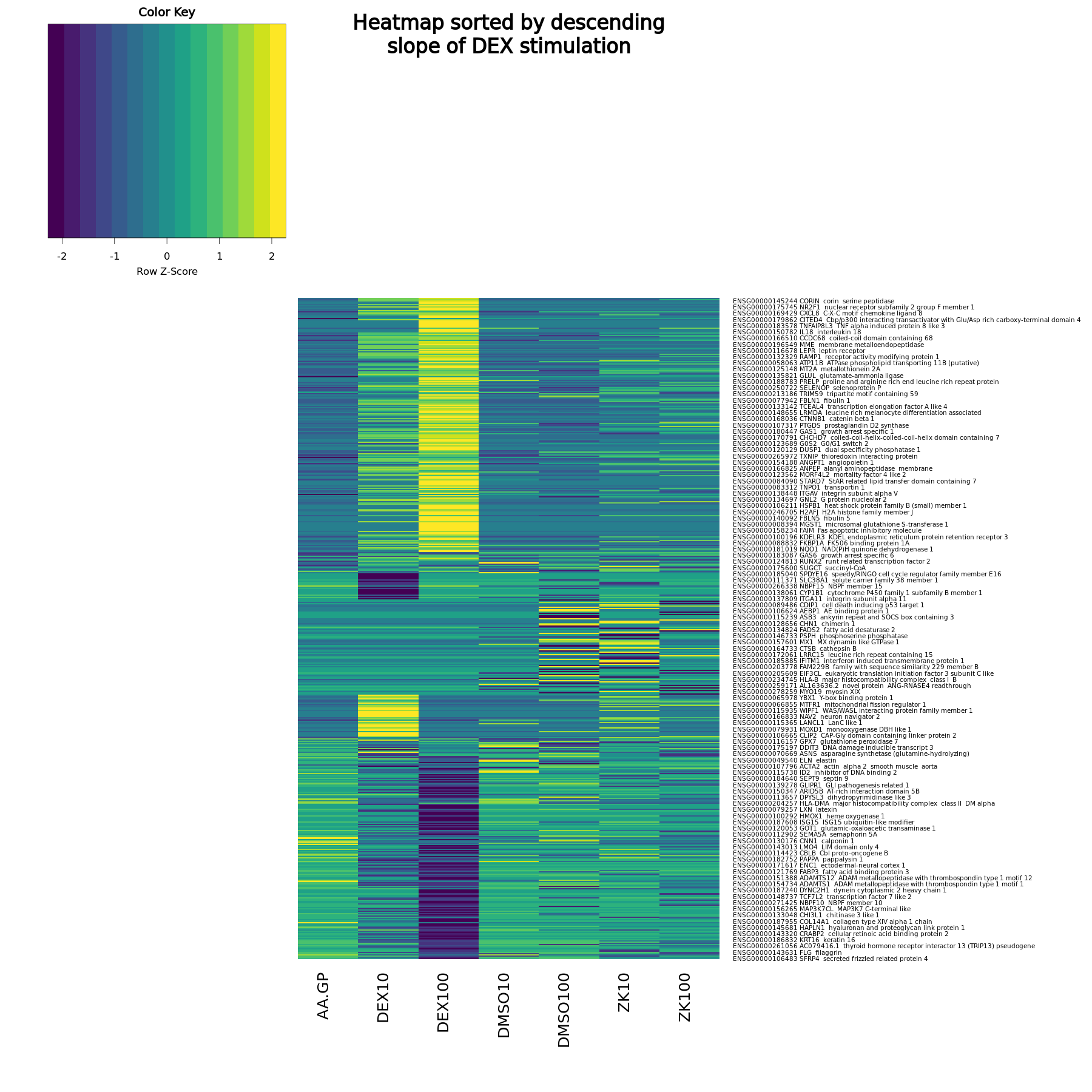
